# Targeted Proteomics of Plasma Extracellular Vesicles Uncovers MUC1 as Combinatorial Biomarker for the Early Detection of High-grade Serous Ovarian Cancer

**DOI:** 10.1101/2022.03.31.486596

**Authors:** Tyler T. Cooper, Dylan Z. Dieters-Castator, Jiahui Liu, Gabrielle M. Siegers, Desmond Pink, Lorena Veliz, John D. Lewis, François Lagugné-Labarthet, Yangxin Fu, Helen Steed, Gilles A. Lajoie, Lynne-Marie Postovit

**Affiliations:** Department of Biochemistry, Western University, London ON, Canada; Department of Biomedical and Molecular Sciences, Queen’s University, Kingston ON, Canada; Department of Anatomy and Cell Biology, Western University, London, ON, Canada; Department of Oncology, University of Alberta, Edmonton AB, Canada; Department of Chemistry, Western University, London, ON, Canada; 6Department of Obstetrics and Gynecology, University of Alberta, Edmonton AB, Canada

**Keywords:** High-grade Serous Carcinoma, Extracellular Vesicles, Proteomics, Biomarkers, Ovarian Cancer

## Abstract

**Purpose:** The five-year prognosis for patients with late-stage high-grade serous carcinoma (HGSC) remains dismal, underscoring the critical need for identifying early-stage biomarkers. This study explores the potential of extracellular vesicles (EVs) circulating in blood, which are believed to harbor proteomic cargo reflective of the HGSC microenvironment, as a source for biomarker discovery.

**Experimental Design:** We conducted a comprehensive proteomic profiling of EVs isolated from blood plasma, ascites, and cell lines of patients, employing both data-dependent (DDA) and data-independent acquisition (DIA) methods to construct a spectral library tailored for targeted proteomics. Our investigation aimed at uncovering novel biomarkers for the early detection of HGSC by comparing the proteomic signatures of EVs from women with HGSC to those with benign gynecological conditions. The initial cohort, comprising 9-10 donors, utilized DDA proteomics for spectral library development. The subsequent cohort, involving 30 HGSC patients and 30 control subjects, employed DIA proteomics for a similar purpose. Support vector machine (SVM) classification was applied in both cohorts to identify combinatorial biomarkers with high specificity and sensitivity (ROC-AUC > 0.90).

**Results:** Notably, MUC1 emerged as a significant biomarker in both cohorts when used in combination with additional biomarkers. Validation through an ELISA assay on a subset of benign (n=18), Stage I (n=9), and stage II (n=9) plasma samples corroborated the diagnostic utility of MUC1 in the early-stage detection of HGSC.

**Conclusions:** This study highlights the value of EV-based proteomic analysis in the discovery of combinatorial biomarkers for early ovarian cancer detection.

## 1. Introduction

Despite an increasing understanding of epithelial ovarian cancer (EOC) etiology and biology, EOC remains the most lethal gynecological cancer in developed countries^1^. Globally, approximately 200,000 women are diagnosed per year, with a 5-year survival rate that remains below 50%^2^. Early detection of HGSC is crucial to improving outcomes, with 92% of patients surviving following early-stage detection, versus only 29% in late-stage cases. Unfortunately, 75% of women experience non-specific symptoms (e.g. abdominal discomfort) and are not diagnosed until the disease has progressed to stage 3 and beyond. In many cases these non-specific symptoms lead to the identification of pelvic masses by transvaginal ultrasound (TVUS) imaging. If abnormal masses are identified, invasive surgical procedures, tissue debulking, and pathohistological analyses are then required to discriminate between benign and malignant disease. High-grade serous carcinoma (HGSC) is the most lethal and aggressive form of epithelial ovarian cancer, accounting for >75% of EOC cases. The extracellular epitope of MUC16 (CA-125) can be used to monitor the progression of EOC and response to chemotherapeutics in combination with TVUS^3, 4^. Unfortunately, tests for CA-125 are not sensitive nor specific enough for early diagnosis of malignant EOC^3^. For example, although ∼20% of patients with late-stage EOC exhibited elevated CA-125 levels (>35 U/mL), increased CA-125 was also observed in women with alternative gynecological conditions^5, 6^. Thus, there remains a dire need to discover alternative biomarkers to aid in the early detection of HGSC.

Algorithms, such as the Risk of Malignancy Index (RMI), aim to incorporate menopausal status, CA-125 levels and TVUS imaging. Alternatively, the Risk of Ovarian Cancer Algorithm (ROCA) monitors CA-125 levels over time to assess the risk of developing ovarian cancer. Unfortunately, large randomized control trials (US Prostate, Lung, Colorectal and Ovarian Cancer Screening Trial and UK Collaborative Trial of Ovarian Cancer Screening) involving thousands of women found no significant survival benefit for multimodal screening strategies over standard of care^5, 6^. Alternative biomarkers to CA-125 have been proposed for estimating HGSC risk. For example, the risk of ovarian malignancy algorithm (ROMA) monitors human epididymis protein 4 (HE4 or WFDC2) in addition to CA-125^6^. The FDA-approved OVA1 in vitro diagnostic multivariate index assay measures five biomarkers (CA-125, transferrin [TF], transthyretin (prealbumin), apolipoprotein A1 [APOA1], and beta-2 microglobulin [B2M]) and demonstrates improved prediction accuracy of malignancy risk compared to a physician’s pre-operative assessment or CA-125 alone^7^. Moreover, Yip *et al.* screened 259 serum biomarkers from HGSC patients and identified nine combinatorial biomarkers with greater specificity than OVA1 (88.9 versus 63.4%)^8^. Høgdall *et al.* screened serum from 150 HGSC patients and found B2M, TF, and ITIH4 robustly predicted overall survival and progression-free survival^9^. These approaches improve cancer classification and monitoring strategies; however, viable biomarkers that can detect early-stage HGSC are still unavailable.

Blood plasma remains an ideal source for biomarker discovery due to the easy acquisition of patient samples for high-throughput immunoassays. Mass spectrometry (MS)-based proteomics is a medium-throughput technique for biomarker discovery; however, the detection of low abundance proteins in plasma is technically complicated by the presence of high abundance proteins (HAPs)^10-14^. Keshishian *et al.* detected ∼5300 plasma proteins by depleting the 14 most abundant plasma proteins as well as ∼50 moderately abundant proteins in tandem with peptide fractionation. Alternatively, N-glycopeptide enrichment can be used to identify plasma proteins relevant to ovarian cancer relapse^15^. It remains to be determined what the optimal strategy is for segregating biomarkers from HAP in primary tissue samples. Extracellular vesicles (EVs), 30-1000nm in diameter, carry bioactive lipid, nucleic acid and proteomic cargo in a lipid membrane that allows for transport through systemic circulation to distant tissues^16^. EVs carry bioactive cargo from or towards a metastatic cancer microenvironment^17^, thus enrichment of EVs may segregate potential biomarkers from HAPs or other liable proteins^18^. A limited number of investigations have attempted to characterize HGSC-EV proteomes using EVs from biofluids^19^.

In our investigation, we adopted a two-pronged approach, utilizing both data-dependent acquisition (DDA) and data-independent acquisition (DIA) proteomics, to meticulously profile the proteome of extracellular vesicles (EVs) derived from two distinct cohorts. The first cohort, comprising 9-10 donors, focused on building a spectral library through DDA proteomics for targeted analysis. In contrast, the second cohort, involving 30 patients with HGSC and 30 control subjects, leveraged DIA proteomics for the same purpose. Our analysis utilized support vector machine (SVM) classification to discern potential biomarkers, leading to the identification of MUC1 as a valuable combinatorial biomarker across both cohorts. The diagnostic potential of MUC1 was assessed using ELISA quantification. This comprehensive approach highlights the efficacy of targeted proteomics and underscores the significance of MUC1 as a biomarker in the early detection of high-grade serous carcinoma.

## Results

### Integrative Proteomic Analysis of Ovarian Cancer Extracellular Vesicles

We first sought to characterize EV proteomes from cancer cell lines, plasma, and ascites fluid for prospective biomarker discovery of early-stage HGSC. Established (OVCAR3, OV-90) cell lines were used to model HGSC, and a non-malignant ovarian surface epithelial cell line (hIOSE) was also analysed. Cell lines derived from ascites fluid of patients with low-grade serous (EOC18) and high-grade (EOC6) ovarian cancer and were also used to reflect a component of the EV proteome generated within an ascites microenvironment. EVs were primarily obtained by UC; however, CD9 affinity purification (CD9AP) was also performed on plasma and ascites to enrich for a subset of EVs (**Figure 1A, S1A**). SCX fractionation was employed to increase proteomic depth prior to LC-MS/MS; in return, >8000 proteins were identified in total. Similar to proteomic analyses of ovarian cancer cell lysates^20^ and Raman spectroscopy characterization^21^, cell line derived EVs harboured unique cargo compared to each other but primary and established cell lines (including hIOSE) clustered along principal components **(Fig 1B-C, Fig S1B**). Importantly, the proteomes of all samples were significantly associated with GO Cellular Component (GOCC) annotations indicative of EV-enrichment **(Fig S1C)**. Cell EV proteomes displayed a 35-45% overlap with Vesiclepedia filtered for EOC cell lines and 65% overlap with Vesiclepedia filtered for ascites fluid (**Fig 1D, S1D)**. Proteomes of cell EVs shared 38-58% of proteins with UC ascites EVs when analyzed individually; however, >84% of proteins detected in ascites EVs were identified across the cell EV proteome **(Fig S1E)**. Compared to UC, CD9AP provided a modest increase in shared proteome coverage between cells and biofluids **(Fig S1F)**. Common proteins were associated with neutrophil degranulation and adaptive immunity **(Fig 1E)**. 2121 proteins detected in cell EVs overlapped with ascites EVs and were associated with adaptive immunity and members of the PDGFB, CXCR and VEGF signalling pathways **(Fig S1G)**. These results support the speculation that the proteomic ‘fingerprint’ of cell EVs may reflect a cross-section of EVs produced within a tumour microenvironment.

**Figure 1.**
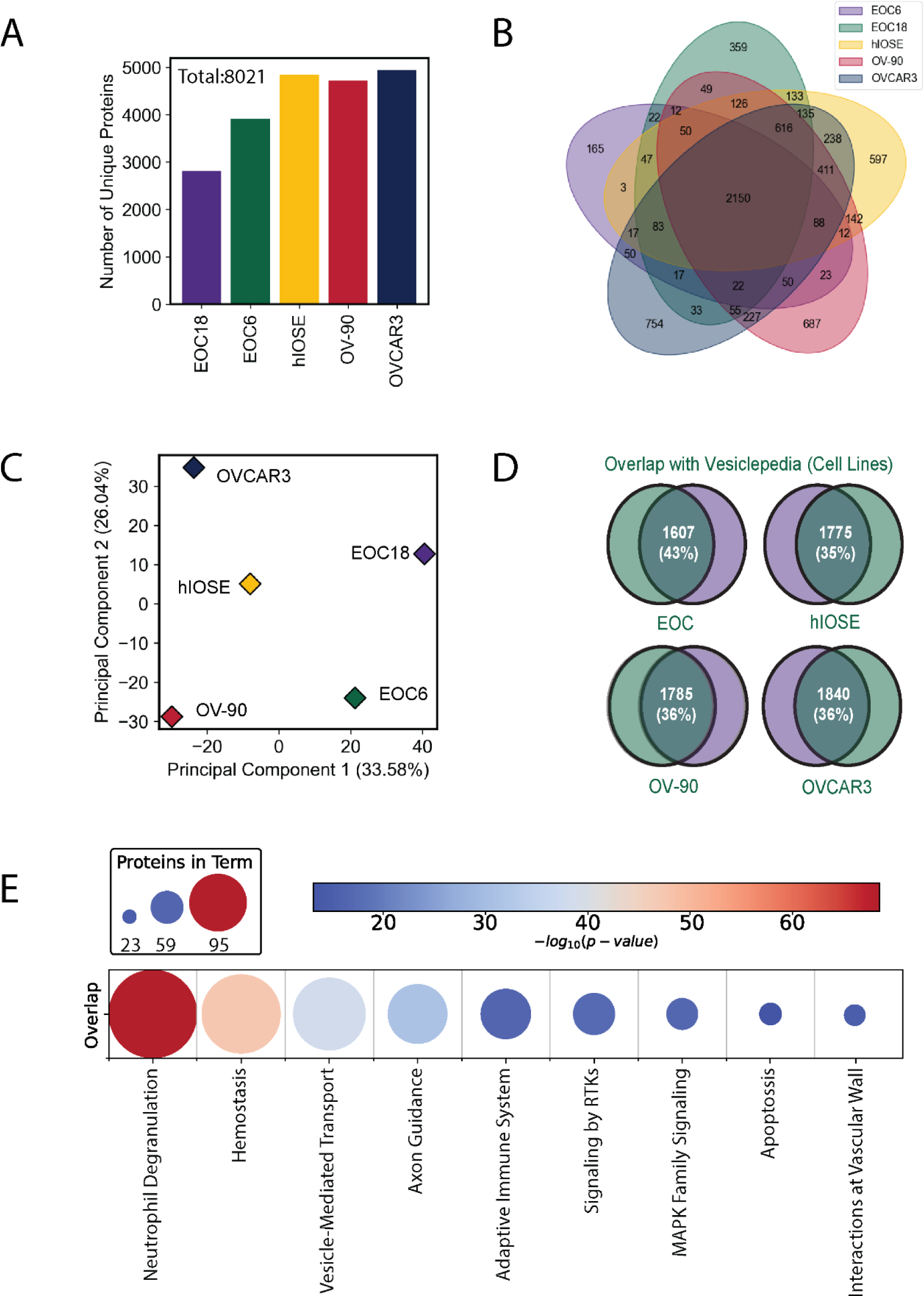
Proteomic Profiling of Extracellular Vesicles Isolated from Ovarian Cancer Cell Lines. EVs were enriched using UC from conditioned media of established (hIOSE, OV-90, OVCAR3) or primary cell lines established from ascites (EOC6 and EOC18). EV proteomes were characterized using UPLC-MS/MS using SCX-DDA. **(A)** The number of unique proteins identified was elevated in EVs derived from established cell lines compared to primary cell lines. **(B)** Venn diagram demonstrating distribution of shared and unique proteins across cell lines. 2150 proteins were identified in EVs from all cell lines. **(C)** Principal component analysis illustrating distinct proteomic ‘fingerprints’ within cell line-derived EVs **(D)** Overlap of EV-proteomes compared to Vesiclepedia database filtered for ovarian cancer cell lines. **(E)** Heatmap of Reactome terms significantly associated with EV proteomes common between cell lines.

### CD9AP increases EV specificity in ascites samples at the expense of proteomic depth

EVs represent a large range of biological vesicles that may reflect anything from ‘cellular debris’ during apoptotic processes to systematically packaged messages that are able to prime distant microenvironments for cancer metastasis^17^. With these properties in mind, we hypothesized that increasing EV purity would uncover additional biomarkers undetected within UC-enriched EV preparations due to EV heterogeneity or residual HAPs. We selected CD9 affinity purification (AP) to segregate a subset of EVs from large EVs and residual cellular debris in ascites. Indeed, EV specificity was increased with CD9AP compared to UC **(Fig 2A, Table S3)**; however, this occurred at the expense of proteomic depth **(Fig 2B,C)**. 145 proteins were exclusively detected in CD9AP-EVs and were enriched with effectors of blood vessel and cancer development, such as TGFB1, BMP2, VEGFC and WNT11. Of note, only PARP1 in CD9AP-EVs overlapped with Vesiclepedia-Ascites **(Fig S2A)**. On the other hand, >1900 additional proteins were exclusively detected using UC of which 1398 proteins were unreported to Vesiclepedia-Ascites **(Fig 2D, Fig S2A)**. Notable mediators of cancer biology exclusively detected in UC-EVs included epidermal growth factor receptor and several components of a innate and adaptive immune response **(Fig S2B)**. 416 proteins were common between isolation methods, although 150 of these proteins were detected at different levels in paired ascites samples **(Fig 2E)**. For example, CD9AP-EVs were enriched for 64 proteins such as tissue plasminogen activator (PLAT) and angiopoietin-like 6 (ANGPTL6). Alternatively, UC-EVs were enriched with 84 proteins, such as Annexin1/2 (ANXA1/2) and myosin heavy chain-9 (MYH9). Although both UC-and CD9AP-EVs contain proteins associated with EV biology, several ‘classical’ EV markers (i.e. CD63) were exclusive to UC-EVs **(Fig S2C,D)**. These results were not surprising considering patterns of CD9 and C63 localization represents distinct mechanisms of EV biogenesis^22^. Integrins facilitate EV uptake into recipient cells and 9 out of 11 detected integrins were exclusively detected in UC-EVs, supporting the enrichment of EV primarily derived from the plasma membrane. This data supports previous reports that increased EV purity with CD9AP can increase the number of prospective biomarkers^23^, albeit putative biomarkers may be lost during this process.

**Figure 2.**
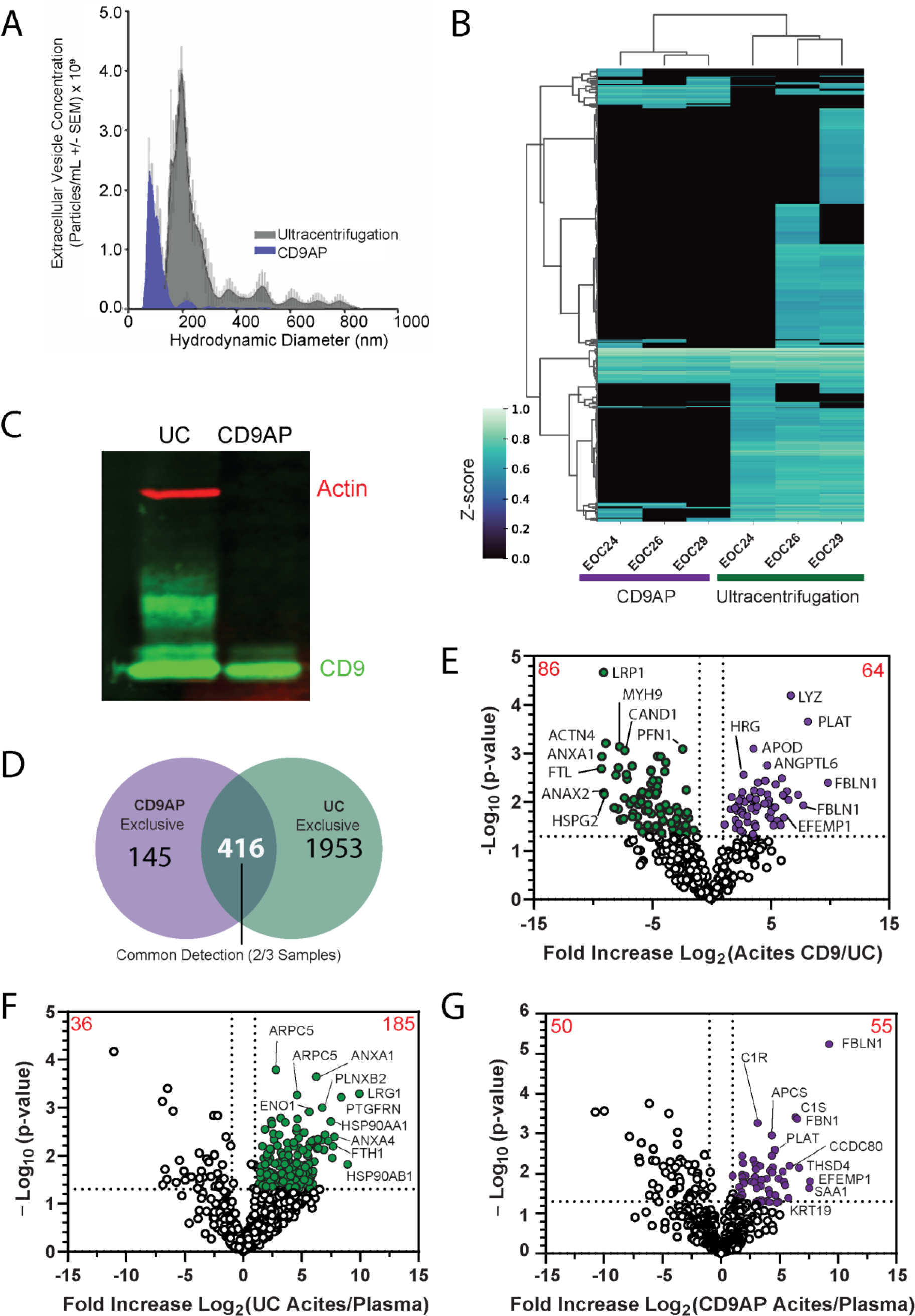
Proteomic Comparison of Ultracentrifugation versus CD9 Affinity purification for Isolation of Plasma or Ascites Extracellular Vesicles. **(A)** Nanoparticle tracking analysis of ascites EVs purified by UC or CD9AP demonstrates a subset of EVs are enriched by CD9AP. The distribution of CD9AP-EVs were primarily distributed around <150nm in diameter, whereas UC samples were comprised of a heterogenous mixture of EVs that were primarily distributed around ∼200nm, albeit subpopulations of EVs were detectable up to 900nm (see Table S3). **(B)** Heatmap of identified proteins and dendrogram demonstrate increased proteomic depth obtained by UC compared to CD9AP using paired ascites donors. **(C)** Cellular debris and large EV components, such as actin, were depleted using CD9AP. **(D)** 145 and 1953 proteins were exclusive to EVs enriched by CD9AP or UC (>1 replicate), whereas 416 proteins were common to both CD9AP and UC-enriched EVs (>2 replicates in each condition). **(E)** Volcano plot of common proteins to CD9AP and UC identified 64 and 84 proteins significantly enriched in either CD9AP- or UC-enriched ascites EVs, respectively. **(F)** 181 proteins were significant enriched within ascites EVs compared to blood plasma EVs collected by UC from healthy donors. **(G)** 55 proteins were significantly enriched using CD9AP on ascites EVs compared to blood plasma EVs collected from healthy donors.

### Quantitative proteomics unveils a large reservoir of putative biomarkers in biofluids

We speculated that ascites EVs derived from the tumour microenvironment would harbour proteomic cargo which may be useful for the detection of HGSC and that a subset of these proteins may be detected within systemic circulation. Accordingly, MS/MS was used to identify proteins enriched in ascites EVs as compared to EVs derived from the plasma of healthy donors. Moreover, if absent in healthy controls, these proteins may be specifically detected in HGSC patients even when tumor burden is low. Like ascites, we employed parallel purification strategies, UC and CD9AP, to increase proteomic depth for biomarker discovery in blood plasma. In the UC group, 185 proteins were significantly elevated (2-fold, p<0.05) in ascites compared to healthy plasma **(Fig 2F)**. These included proteins associated with cancer cell biology and/or metastasis, such as LRP1, and MUC1. On the other hand, 55 differentially expressed proteins (2-fold, p<0.05) were detected between healthy plasma and ascites using CD9AP **(Fig 2G)**. These included cancer-relevant proteins such as MMP14 and CD14. Next, we sought to determine whether ascites-specific EV proteins could also be detected in the plasma of HGSC patients. Over 200 proteins that were enriched within ascites were also detected in plasma samples from donors with HGSC and included mediators of immune response and regulated exocytosis **(Fig S2C, E)**. These proteins were considered as prospective biomarkers during PRM method development. HE4 was not detected in our study, which suggested potential EV- independence, similar to that reported in Zhao *et al^24^*. Collectively, these results support the parallel application of UC and CD9AP to ‘mine’ prospective biomarkers; moreover, confirm that ascites may be a resourceful biofluid for early biomarker discovery.

### Targeted proteomics of plasma EVs and support vector machines identified several biomarker combinations for the early detection of HGSC

A large number of proteins were significantly enriched in ascites EVs as compared to healthy plasma EVs and could be detected in plasma EVs of HGSC patients. Hence, we next asked whether protein abundance could be used to differentiate plasma EV samples from patients diagnosed with HGSC (n=10) versus controls with non-cancerous gynaecological conditions (n=9). Donors were age-matched and ranged from 39-69 years old and an average age of 54.8 **(Table S4).** We chose patients with non-cancerous gynaecological conditions to serve as our controls to account for proteins that may be associated with pathologies or general inflammation in addition to ovarian-cancer-specific analytes which may be released within the tumour microenvironment. A curated list of 471 peptides (240 proteins with evidence in ascites and plasma) was subsequently targeted using a PRM method built in PEAKS^25^ and Skyline^26^ **(Fig S3)**. Peak areas were normalized to the TIC to correct for technical variability, and additionally normalized to the CD9 peptide EVQEFYK (extracellular region, AAs 120-126) to control for EV purity. A total of 21 peptides were significantly different in malignant versus non-malignant samples and were used in further analyses (Wilcoxon rank-sum test, p<0.05) (**Fig 3A**). Of note, a peptide from CA-125 (MUC16) (ELGPYTLDR) was included in our PRM method (ELGPYTLDR). Using the Wilcoxon rank-sum test, this peptide achieved p=0.060 for an AUC of 0.76 and log2 fold-change 2.12. While this did not pass our statistical threshold, we included it in further analyses based on the use of CA125 as a biomarker for HGSC. Based on these 22 peptides, malignant and non-malignant samples were partially segregated using PCA and unsupervised k-means classification (**Fig 3B**). Machine learning classification models, such as SVMs, have demonstrated immense utility for identifying novel biomarkers for an array of diseases. This is due to their ability to provide high-accuracy classification using high-dimensionality data when sample numbers are limited. Indeed, this is an extremely beneficial and attractive feature of SVMs for biomarker discovery studies where the acquisition of large donor numbers is extremely difficult or impossible to obtain. Data features were scaled using z-scores, and randomly split into 10 independent training (70%) and test (30%) sets in a stratified manner. Donor status, such as FIGO stage, remained blinded until final validations were performed using the test set. As proof-of-principle, we retrospectively chose random_state=6 which contained all FIGO stages in both training and test data sets, thus allowing us to speculate on the ability of prospective biomarkers to identify early-stage HGSC. The optimal hyperparameter(s) were determined by LOOCV to reduce variance often obtained with low complexity data sets by reserving a single sample for validation^27^**(Fig S3)**. 14,784 total fits or permutations were used to calculate a mean accuracy score using Matthew’s Correlation Coefficient. From these analyses, we identified eight linear SVMs (C=0.025-2) that provided a mean accuracy score >90% **(Fig 3C)**. Next, we optimized feature selection based on Receiver Operating Characteristic-Area Under the Curve (ROC-AUC) using the reserved test set. The SVM (PC=2, C=0.025) was tested 231 times with paired permutations of all 22 peptides (Appendix Fig 5). Interestingly, nine combinations of peptides were able to classify malignant (n=4) versus non-malignant (n=3) samples with a ROC-AUC = 1.0 **(Figure S4A)**. For example, the combination of CFHR4 and MUC1 was able to accurately classify Stage I, II, and III donors in comparison to MUC16 **(Fig 3D-F)**. Several peptide combinations provided ROC-AUC = 1.0, however GPX3, MUC1, and CFHR4 were represented in the majority of models **(Table S5)** CFHR4 and GPX3 were not detected in cell line EVs and MUC1 was not detected within CD9AP-EVs; yet, all were considered strong drivers of HGSC classification according to SHapley Additive exPlanations (SHAP) analysis^28^**(Fig S4B,C)**. Interestingly, CFHR4 was also considered a strong driver of SVM accuracy in EV-depleted plasma **(Fig S5)** and was speculated to be constituent of the EV corona^29^. Ultimately, we highlight the use of label-free PRM, SVM optimization using LOOCV and parallel enrichment of EVs to identify combinatorial biomarkers of HGSC.

**Figure 3.**
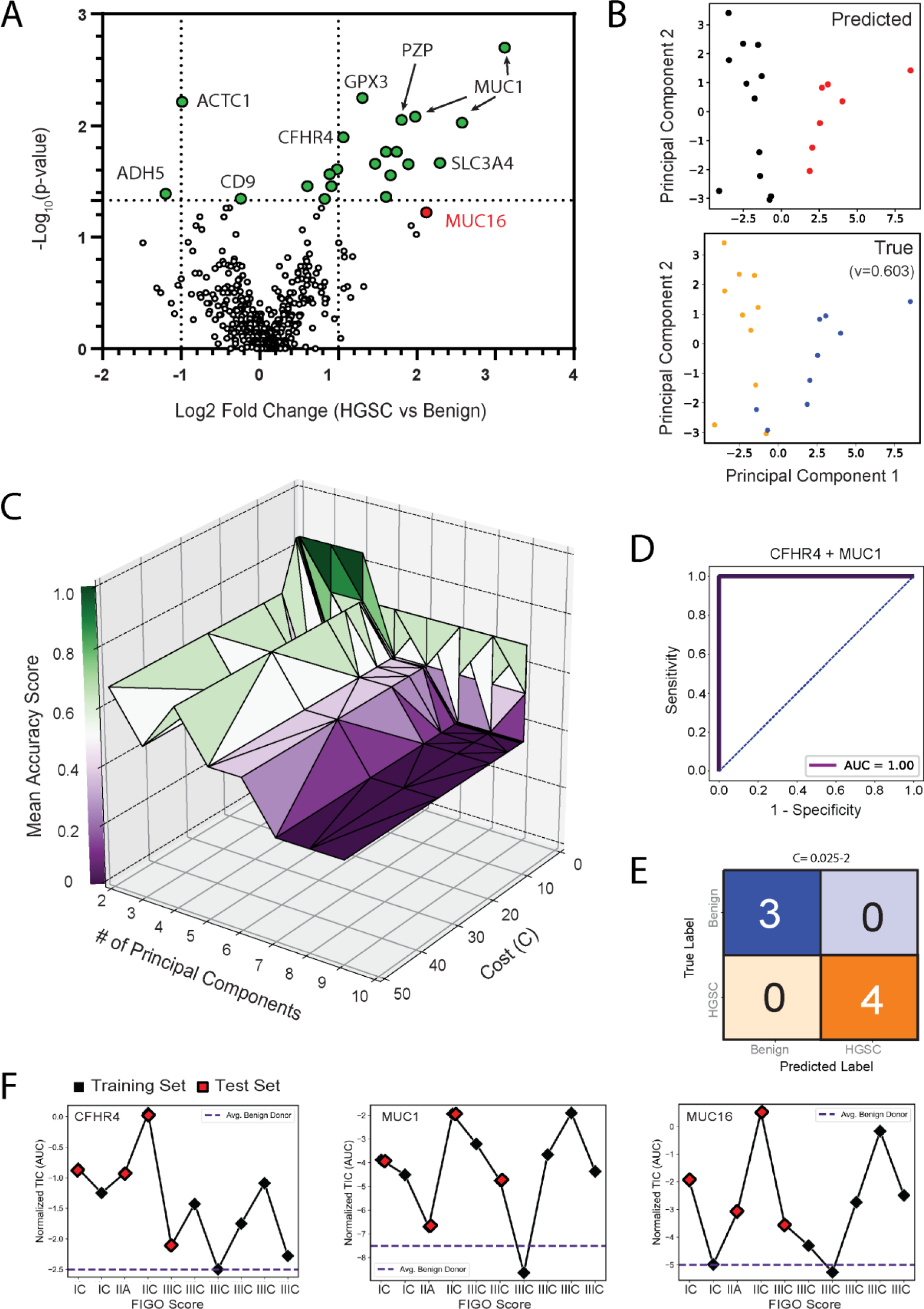
Targeted Proteomics and Support Vector Machine Classification Identifies Prospective Biomarkers to Distinguish HGSC vs Benign Disease. 471 peptides corresponding to 240 proteins were analysed in EV-enriched blood plasma from HGSC (n=10) versus control (n=9) donors using PRM. **(A)** Volcano plot highlights peptides that were significantly different between malignant and control donor samples. 21 peptides (p-value <0.05) and HGSC antigen MUC16 (red) were selected for further analyses. **(B)** Unsupervised PCA and k-means clustering of pooled samples. Predicted labels (red and black) partially overlapped with true labels (blue = Benign and orange = HGSC). V-measure = 0.603. **(C)** Hyperparameter tuning of the linear SVM was performed by LOOCV, leading to hyperparameters C=0.025-1 and two principal components selected as the ‘optimized’ SVM based on mean accuracy score (>0.90). Each point of triangulation indicates an SVM combination/fit that was scored using the training set. Feature selection was performed using 231 combinations of peptides and test data. From this analysis, nine combinations of peptides provided an accuracy score of 1.0 on the test data set (see Figure S4A). **(D-E)** For example, the combination of CFHR4 and MUC1 provided a Receiver Operating Characteristic-Area Under the Curve (ROC-AUC) score of 1.0. **(F)** Training (red) and test samples (white) were represented by women with Stage I, II, and III EOC.

### Size Exclusion Chromatography and Data-Independent Library Generation Uncovers MUC1 as a Prospective Biomarker using Targeted Proteomics

We next focused our biomarker discovery pipeline to a larger cohort of patients diagnosed with FIGO I/II HGSC (n=30) versus controls with non-cancerous gynaecological conditions (n=30). Donors were age-matched and ranged from 40-82 years old with an average age of 62.8 **(Table S6, Figure S6).** In these analyses aiming to identify early-stage biomarkers, we opted to utilize size-exclusion chromatography to increase the purity of plasma EVs while retaining sufficient yield for MS analyses. Indeed, EV concentration and size profiles using nanoparticle tracking analysis **(Fig 4A-B)** and atomic force microscopy align **(Fig 4C-D)** with previous reports of plasma EVs. Instead of offline peptide fractionation (i.e., SCX), we opted to implement a gas-phase fractionation technique for increasing proteomic depth during DIA-based biomarker discovery. In this approach a collection of DIA in 100 m/z windows were acquired across 300-1000 m/z, thus equalling 7 injections with 2 m/z isolation windows after demultiplexing MS/MS spectra. Next DIA acquisitions between 400-1000 m/z were acquired on pooled EVs isolates to build a spectral library of detectable peptides for targeted analyses. Using this approach, we were able to identify >2000 proteins across plasma EVs of which >1800 contained a unique peptide. Aligned with UC EVs, SEC EVs were enriched with proteins associated with neutrophil degranulation, wound healing, and blood microparticles **(Fig 4E).** We selected peptides that provided reproducible retention times in comparison to internal heavy isotope standards (r > 0.95). Additional manual curation removed high abundant proteins, such as albumin, that would likely provide little diagnostic value. In total, we focused on 290 proteins totaling 495 peptides across two independent PRM methods. PCA of our targeted proteomics highlighted the similarity of early-stage HGSC vs benign gynecological disease; albeit centroids of HGSC and Benign donors were separated on principal components that coincided with differential expression levels between the two groups **(Fig 5A-B)**. SERPIND1, AGT, and PROZ were enriched within EVs from benign donors, whereas peptides for MUC1 and CD9 were significantly elevated in HGSC EVs **(Fig 5B)**. Peptides for MUC1, ‘QGGFLGLSNIK’ and ‘DISEMFLQIYK’, were elevated in HGSC EVs 3.14- and 8.86-fold over benign EVs, respectively. Interestingly, CD9 peptide (TKDEPQRETLK) was elevated 4.43-fold in HGSC EVs and was the large contribution to principal component 1. Unfortunately, our NTA analysis did not assess if elevated CD9 corresponded to an increased number of CD9 particles between HGSC versus benign controls. Similar to our first cohort, the level of MUC16/CA-125 was not found to be significant compared to benign controls.

**Figure 4.**
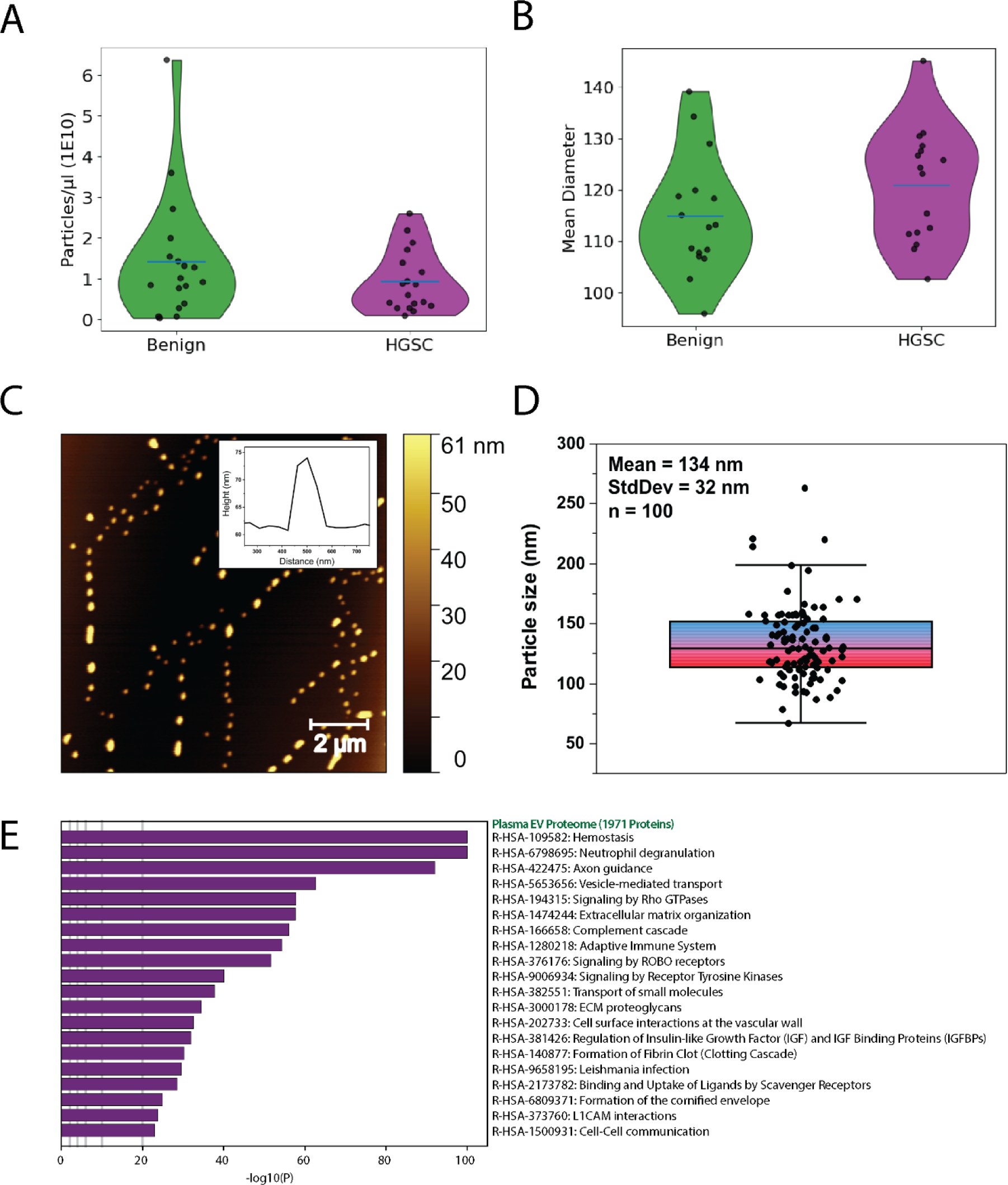
Isolation of Plasma EVs by Size Exclusion Chromatography. EVs were isolated using size-exclusion chromatography from plasma of patients with early-stage HGSC (n=16) and benign gynecological disease (n=16). EV concentration and mean diameter were estimated using nanoparticle tracking analysis. EV isolations from HGSC and Benign donors displayed comparable **(A)** particle concentrations and **(B)** average mean diameter of particles detected. Blue horizontal lines represent mean. NTA estimated a mean diameter of 120µm for plasma EVs. **(C-D)** Topographical analysis of HGSC EVs by atomic force microscopy detected EV-like particles with mean diameter of 134nm. **(C)** Insert a representative line trace of a single particle, displaying a dome like structure. **(D)** Data represented as mean and standard deviation using box plot with inner quartiles shown. **(E)** Enrichment Analyses of Reactome and GO Biological Process annotations in comparison to total proteins identified by GPF-DIA MS/MS.

**Figure 5.**
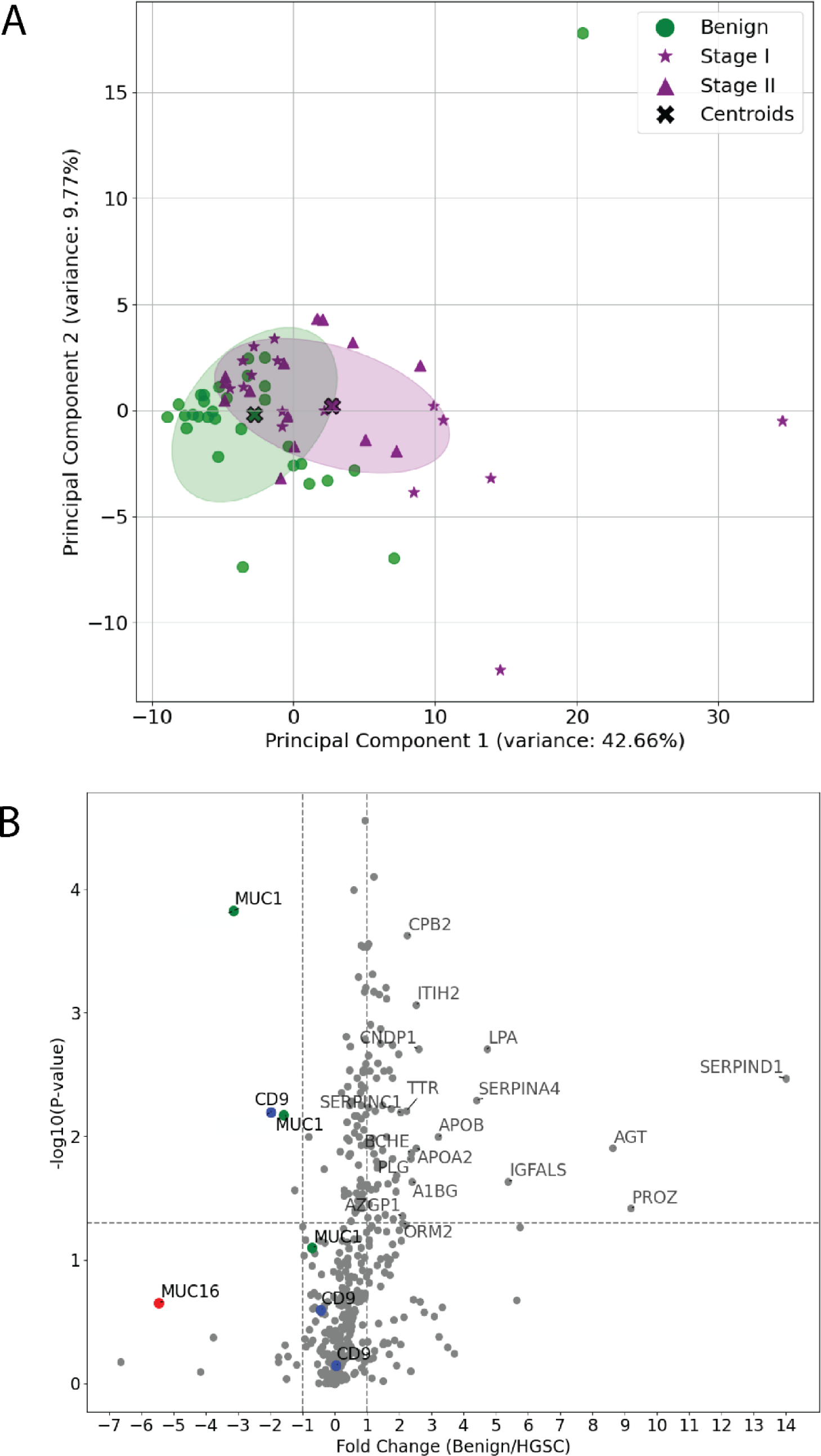
Differential Expression and Principal Component Analysis of PRM Analysis. **(A)** Principal Component Analysis (PCA) of targeted proteomics comparing plasma EVs from early-stage high-grade serous carcinoma (HGSC) patients (stages I and II, n=30) to controls with benign gynecological conditions (n=30). The PCA plot displays the distribution of samples with benign conditions (green circles), stage I HGSC (purple stars), stage II HGSC (purple triangles), and centroids (black crosses) indicating the average position of each group. **(B)** Volcano plot illustrating the differential protein expression between benign and HGSC EVs. X axis indicated log_2_(fold change). Y axis indicate -log10(p-value). Proteins of particular interest, such as MUC1, CD9, and MUC16, are highlighted and labeled.

Implementation of our SVM optimization pipeline found that the accuracy of SVM was increased up to 0.95 with increasing features up to 10, albeit SVM with 2-3 features were able to provide 0.7-0.8 mean accuracy scores on a training set **(Figure 6A)**. In order to keep the SVM simple, we decided to assess all combination of two feature SVM using a range of cost (C) weights. This approached determined C=1.0 was able to identify several combinations of proteins that provided a ROC-AUC >0.85 on a test set **(Figure S7A)**. For example, MUC1 and APOC4 were able to correctly classify 8 out of 10 HGSC donors and all 10 Benign donors, equaling an ROC-AUC = 0.90 **(Figure 6B)**. APOC4 is currently a under review by the FDA as a biomarker for ovarian cancer (https://edrn.nci.nih.gov/data-and-resources/biomarkers/apoc4/) and demonstrated a high ROC-AUC (0.87) using logistic regression within this study **(Figure S7B)**.

**Figure 6.**
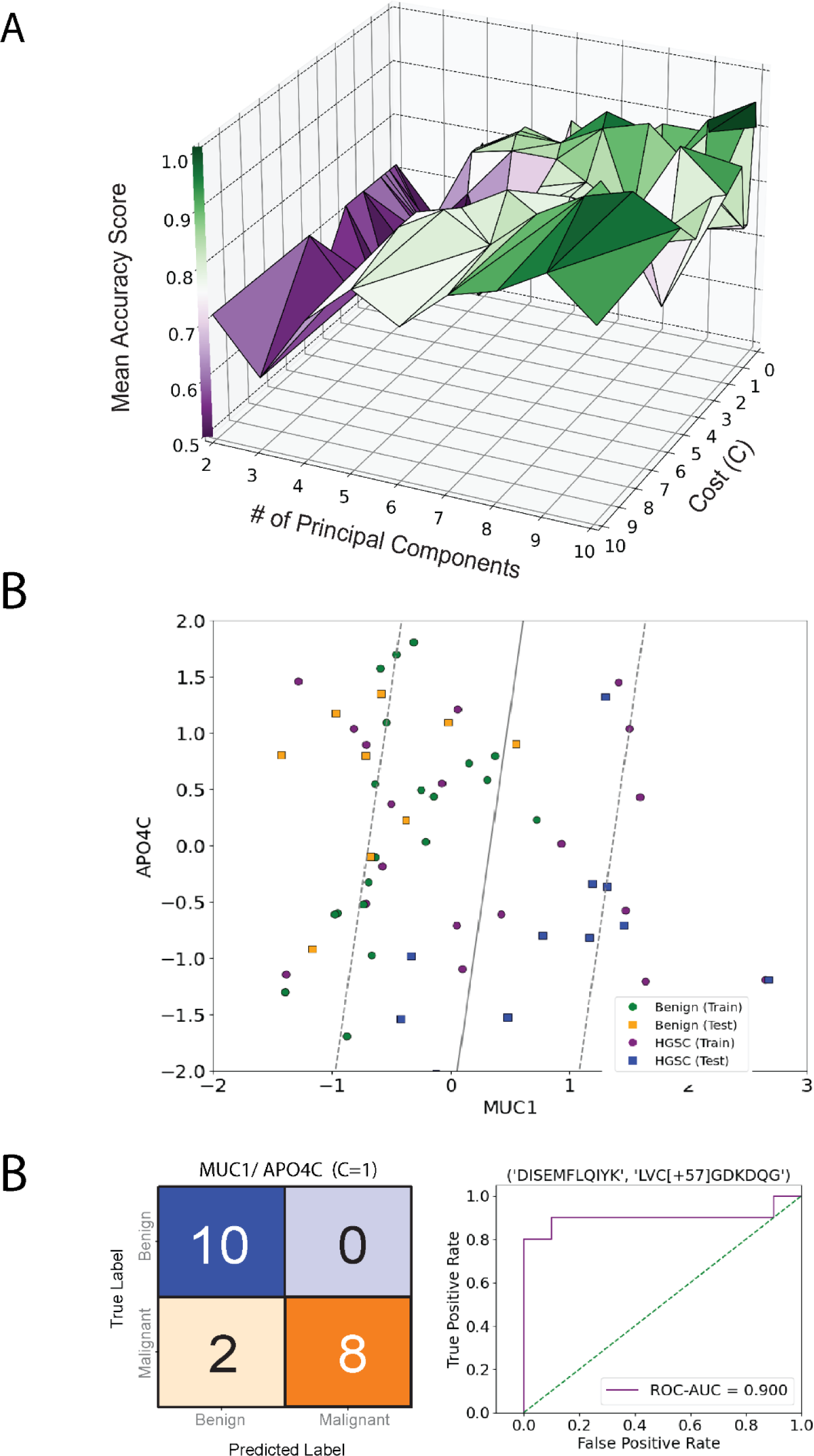
Exploration of Support Vector Machines to Uncover Prospective Biomarkers Capable of Classifying Early-Stage HGSC vs Benign Gynecological Disease. Peptides with p<0.05 were selected as features for support vector machine model training and validation. HGSC and Benign donors were split into training and test data sets. **(A)** SVM training using LOOCV was used to determine optimal cost (C) and number of principal components or features to maximize prediction accuracy determined by Matthew’s Correlation Coefficient or mean accuracy score. Within these analyses, model accuracy was increased with increasing features, however cost weight had less of an influence. Using 2 feature models, we identified several combinatorial peptides which provided high sensitivity and specificity using the test data set. For example, **(B-C)** support vector machine utilizing APOC4 and MUC1 was able accurately classify 8 out of 10 HGSC donors and 10 out of 10 Benign donors. ROC-AUC for this model was determined to be 0.90.

### Plasma MUC1 is a prospective biomarker for the detection of early-stage HGSC and increases with tumour progression

Like SHAP analysis of SVMs trained with the proteome UC EVs, MUC1 was classified as a strong driver of HGSC vs Benign classification in SVM models built on proteome data from SEC-isolated EVs **(Fig 7A)**. Two independent PRM studies identified MUC1 as a prospective biomarker for early-stage HGSC detection. Accordingly, we wanted to determine whether MUC1 levels may be able to predict HGSC occurrence and/or progression. Using an ELISA for MUC1 we estimated that the optimal threshold for HGSC detection was 22.31 mU/mL. Using this threshold, we detected elevated MUC1 in raw plasma samples from HGSC donors as compared to donors with benign disease. **(Fig 7B)**. There was also a significant increase in MUC1 levels between FIGO Stage I and Stage II **(Fig 7C)**. Indeed, HGSC was detected with an ROC-AUC = 0.73 **(Fig 7D,E)**. These results aligned with our PRM analysis in which MUC1 yielded a ROC-AUC = 0.75. Finally, logistic regression of MUC1 levels in FIGO I vs FIGO II donors was able to generate an ROC-AUC = 0.93 **(Fig 7F,G)**. Taken together, we provide evidence that MUC1 is a prospective biomarker that can augment the classification of early-stage HGSC from benign gynecological disease when used in combination with additional biomarkers.

**Figure 7.**
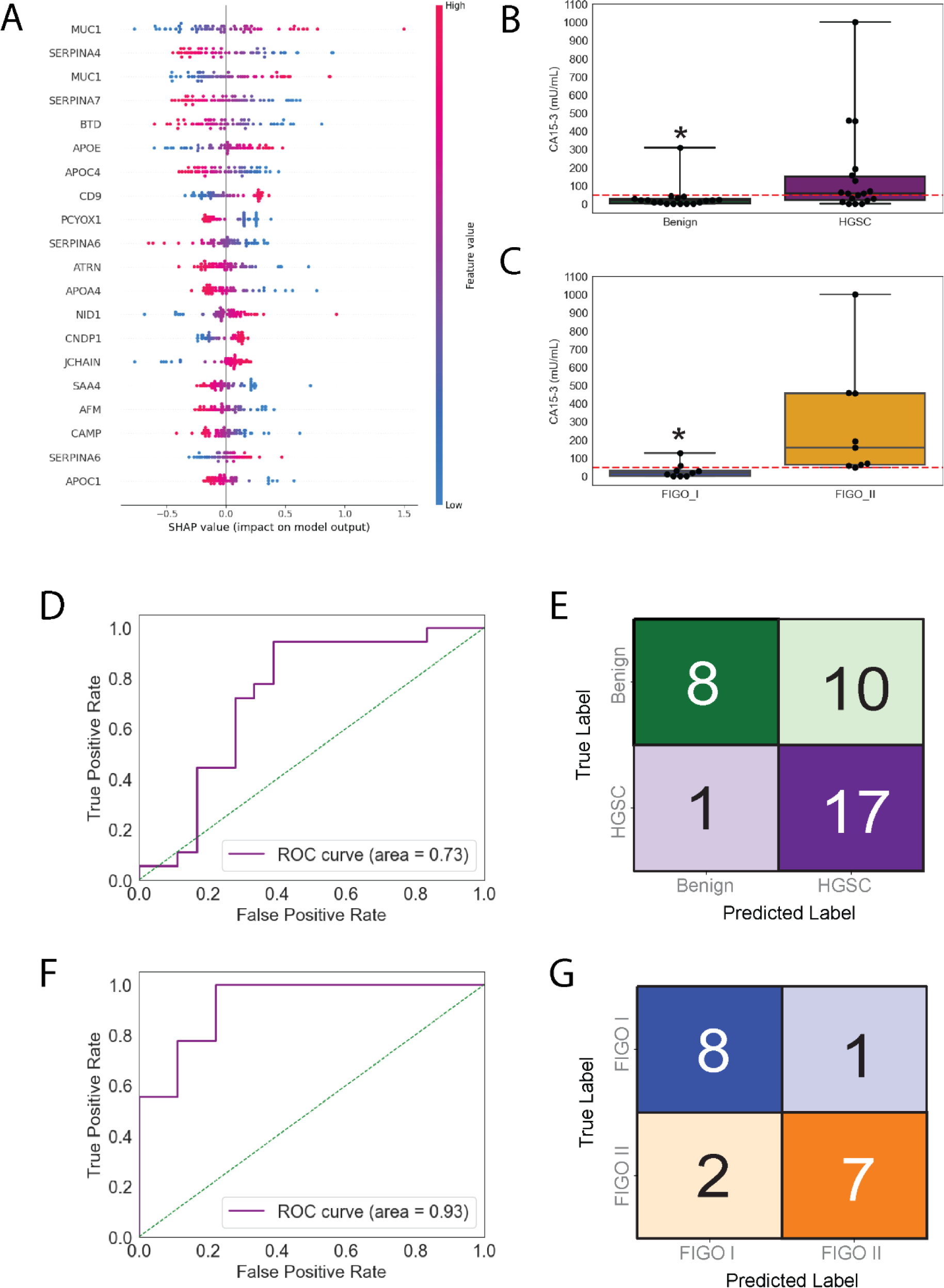
Validation of MUC1 as prospective biomarker for early-stage HGSC. (A) SHAP model analysis reveals strong drivers of SVM classification. Positive SHAP values indicate strong drivers of HGSC classification, such as MUC1. On the other hand, APOC4 was determined as a strong driver of Benign donor classification. MUC1 (CA15-3) concentrations were estimated in raw plasma using ELISA. **(B)** MUC1 was significantly elevated in early-stage HGSC donors compared to donors with benign disease. Notably, **(C)** MUC1 was also differentially detected between HGSC donors classified as FIGO I vs II. **(D-E)** Using logistic regression for classification, MUC1 produced an AUC-ROC = 0.73 and correctly identified 17 out of 18 HGSC donors; albeit 10 out of 18 Benign donors were misclassified. **(F-G)** Logistic regression classification of HGSC donors into FIGO I vs FIGO II provided an AUC-ROC = 0.93. 7 out of 9 HGSC donors at FGIO II were correctly classified; moreover, 8 out of 9 HGSC donors were correctly classified as FIGO I. Data in B,C represented a box plots with quartiles. Red dashed line indicates predicted threshold of MUC1 to confidently indicate HGSC vs benign disease. * = 0.05 > p-value determined by Mann-Whitney U Test.

## Discussion

In this study, we characterized EV proteomes derived from HGSC cell lines, ascites and plasma using two distinct enrichment strategies (UC and CD9AP) to maximize proteomic depth and increase the number of biomarker candidates. Our findings expand upon previous work by several other groups that also utilized mass spectrometry to characterize EVs derived from cells or biofluids. In stark contrast to the previous studies, we were able to build SVM models using targeted proteomics to identify early-stage HGSC from plasma EVs in comparison to benign disease.

Our comparisons of EV proteomes from cell lines supports previous reports of intercellular heterogeneity, which may reflect differences in tissue of origin or stages of ovarian cancer progression^30^. For example, three distinct proteomic expression profiles were identified during a large-scale proteomic analysis of cell lines and primary tumors^31^. We found the EV proteomes of cell lines may reflect the pathophysiology of early-stage HGSC, such as inflammation^32^, ECM remodeling^33^ and angiogenesis^34^. However, many similarities were noted between cancer cells and the non-malignant hIOSE, pointing to potential confounders associated with propagation in tissue culture. Building off the proteome of HGSC cell line EVs, we expanded our focus to the proteomic profiling of EVs from primary sources. We executed an in-depth characterization of biofluid-EV proteomes using parallel purification strategies, the ‘match-between-runs’ feature in MaxQuant^35^, SCX StageTip fractionation technology^36^, GPF-DIA^37^ and Orbitrap-based instrumentation. Several efforts have attempted to deplete HAPs from biofluids to improve the detection of low-abundancy biomarkers^38, 39^, however a consensus on the optimal method has yet to be determined. To better delineate proteins specific to HGSC, Shender *et al.* compared ascites from patients with ovarian cancer to those with alcohol-induced cirrhosis and identified 424 proteins associated with malignant ascites. More recently, Sinha *et al.* have developed an HGSC xenograft model in combination with N-glycopeptide enrichment and PRM to identity potential biomarker candidates in primary patient samples^40^. Considering the proteomic complexity of biofluids, it is unlikely that a single proteomic approach will be able to identify all biomarkers for detecting metastatic HGSC.

Within this study, we demonstrate a robust pipeline incorporating EV purification, PRM proteomics, and SVM that is tailored for the identification of prospective biomarker combinations for early HGSC detection. In agreement with previous studies, MUC16 was elevated in malignant samples but was not considered a stand-alone biomarker. Combinations of MUC16 and additional peptides were able to provide high accuracy; however, subsequent investigations with larger cohorts will be necessary to determine the diagnostic value of the combinatorial biomarkers uncovered in this study. SHAP analysis can provide additional insight into which peptides drive prediction outcomes within a machine learning classification model^28^. Using these analyses, we identified MUC1 as a strong driver of HGSC classification using EVs purified by both UC and SEC. MUC1 is a single-pass transmembrane protein that is significantly upregulated in HGSC. Furthermore it is subjected to isoform splicing and deglycosylation during tumorigenesis^41^. In the context of cancer, the extracellular domain of MUC1 is cleaved and released into systemic circulation wherein it appears to contribute to several intercellular signaling networks via RTK, EGFR and Akt interactions^42-44^. MUC1 has been proposed as a biomarker for HGSC monitoring and was elevated in HGSC EVs isolated from pooled plasma^45^. This study provides two additional proteomic analyses that identify MUC1 as a prospective biomarker in plasma EVs of HGSC donors. We validated our PRM analyses by employing ELISA quantification on the MUC1 antigen CA15-3. Indeed, MUC1 was significantly elevated in HGSC donors and provided high sensitivity using logistic regression. Unfortunately, MUC1 provided an ROC-AUC = 0.73, thus it cannot be considered as a stand-alone biomarker. It is likely necessary to obtain the sensitivity and specificity for clinical application^6, 9, 46^. Accordingly, we identified several protein combinations with MUC1 that provided high accuracy (ROC-AUC >0.9). Future efforts will need to validate combinatorial biomarkers using an independent cohort of HGSC vs Benign donors. Notably, plasma MUC1 was significantly elevated in FIGO II compared to FIGO I, thus supporting the idea that circulating CA15-3 increases with tumour burden. We did not measure glycosylation levels on MUC1 in plasma EVs from early-stage HGSC, thus future research may benefit from the enrichment of glycopeptides for PRM analyses^40^. Recently, Wenk *et al.* identified a glycopeptide from the protein Latent Transforming Growth Factor Beta Binding Protein 1 (LTBP1) that outperformed MUC16 for monitoring remission and recurrence in a cohort of patients with HGSC^47^. Taken together, additional proteomic techniques may be useful to uncover prospective biomarkers for early-stage detection of HGSC.

A limitation of our study was the absence of platelet depletion in biofluid samples prior to cryopreservation, likely limiting proteomic depth due to an abundancy platelet proteins^48-50^. This issue, common in proteomic research, might have obscured low-abundance, HGSC-specific proteins. Restricted sample volumes obtained from biobanks limited our ability to validate additional biomarkers besides MUC1. Nevertheless, our methodology robustly identified combinatorial biomarkers with high diagnostic specificity and sensitivity for early-stage HGSC. Future studies will need to consider all aspects of plasma collection and storage to enhance our biomarker detection pipeline. Despite this challenge, our findings contribute significantly to early HGSC diagnostics and highlight the potential for early intervention strategies focused on EV proteomes.

## Methodology and Data Analysis

### Cell Culture

OV-90 (ATCC® CRL-11732) and NIH:OVCAR3 (ATCC® HTB-161) were obtained from the ATCC. Human immortalized surface epithelial cells hIOSE (OSE364) were obtained from the Canadian Ovarian Tissue Bank at the BC Cancer Agency and kindly provided by Dr. Ronny Drapkin (Department of Obstetrics and Gynecology, University of Pennsylvania). Primary cell lines EOC6 and EOC18 were isolated from the ascites of patients with high-grade and low-grade serous ovarian cancer, respectively. All cell lines, except OVCAR3, were maintained in M199+MCDB105 supplemented with 5-15% FBS. NIH:OVCAR3 cells were cultured in RPMI-1640 supplemented with 20% FBS and 5µg/mL insulin. Media was exchanged with serum free media for 20-30 hours to generate conditioned media (CM) for EV purification. All work involving the use of patient samples (cell lines, plasma and ascites) was approved by the Health Research Ethics Board of Alberta-Cancer Committee.

### Ascites Fluid

Institutional approval for research with human materials was received prior to the initiation of these studies (Health Research Ethics Board of Alberta-Cancer Committee, HREBA.CC-17-0450), and samples were obtained after receiving informed consent. Briefly, ascites fluid aspirates were depleted of cells and cellular debris through serial centrifugation at 300g for 10mins and 1000g for 10 mins, respectively. 1mL of cell-free ascites fluid was used for each EV isolation. Ascites fluid was stored at -80C.

### Blood Plasma

Blood plasma was collected from treatment naïve-women with HGSC or benign gyneological disease at the University of Alberta after receiving informed consent. Additional plasma samples from women with early-stage (I/II) HGSC or non-cancerous gynecological ailments were obtained from the Banque Cancer de l’ovaire, Centre de recherche du CHUM (CRCHUM), in Montréal, Québéc, Canada. Plasma samples were collected between the years of 2009–2021 from individuals diagnosed with HGSC and before any treatment (chemotherapy or radiotherapy). Plasma from women with HGSC (n = 30) or age-matched controls with benign gynecological ailments (n = 30). Plasma was stored at -80 ⁰C prior to experimentation.

### Ultracentrifugation (UC)

CM, plasma and ascites samples were first centrifuged at 200-300 x g at 4°C to pellet cells. Supernatants were diluted 1:10 in PBS (except CM) and centrifuged at 3,000 x g for 20 minutes at 4°C to remove cell debris. To remove large membrane fragments, supernatants were spun at 10,000 x g for an additional 20 minutes at 4°C. Lastly, supernatants were ultracentrifuged at 120,000 to 140,000 x g (SW-28 rotor) for 2 hours at 4°C to pellet EVs on an OptimaTM L-100 XP ultracentrifuge (Beckman Coulter). The supernatant was removed and EVs were resuspended in 100-300µL of PBS and stored at - 80°C until further use.

### CD9-affinity Purification (CD9AP)

Hydrophilic streptavidin magnetic beads (120mg) were washed three times with PBS then resuspended in 5mL PBS (New England Biosystems, S1421S, 20mg/5ml). Beads were mixed with 650µg biotin conjugated anti-CD9 antibody (Abcam, ab28094) at room temperature for 30 minutes and then washed twice with PBS to remove unbound antibody. Beads were resuspended in 6mL PBS and 1mL (∼20mg) was added to 10mL plasma or ascites (diluted 1:1 in PBS). Samples were placed on a rotary mixer overnight at 4°C and then rinsed three times with PBS. EVs were eluted from beads with three-500 µl glycine-HCl (0.1M, pH 2.39) washes. A small volume (75µL) of Tris-HCl (1.8M, pH 8.54) was used to neutralize each eluent.

### Size Exclusion Chromatography (SEC)

200 µl of benign, or HGSC plasma was loaded onto an Izon 70nm Gen2 column, according to manufactures instructions.. Following three 1.5 mL washes with PBS, ∼200µl experimental plasma is loaded on the column and allowed to enter the column for 5 minutes. Next, 2.0 mL of PBS is loaded into the column and eluent is disposed of until flow from the column is stopped. Finally, 1.5 mL of PBS containing 2.5 mM trehalose was added to SEC columns and up to 1.2 mL was collected and considered EV-enriched. Aliquots were stored at -80 ⁰C until experimental use.

### Western Blotting

EVs were lysed in RIPA buffer. 10 µg protein was loaded onto a 10% SDS-PAGE gel under reducing conditions. Proteins were transferred to PVDF and the membranes were blocked with LI-COR Intercept Blocking solution. Membranes were incubated with anti-CD9 rabbit antibody [CD9 (D8O1A) Rabbit mAb, Cell signaling Tech; #13174S, dilution 1:2000] and an anti-actin mouse antibody [Anti-β-Actin Antibody (C4), Santa Cruz Biotech, sc-47778, dilution 1:1000] overnight at 4°C. Membranes were washed then incubated with IRDye 800CW donkey anti-rabbit (LI-COR# 926-32213, dilution 1:20000) and IRDye- 680RD donkey anti-mouse (LI-COR# 926-68072, dilution 1:20000) for 1 hour at room temperature. Membranes were then scanned with the Odyssey Infrared Imager (LI-COR).

### Nanoparticle Tracking Analysis

Samples were diluted 25-fold using filtered 0.2x phosphate buffered saline and then were analyzed using the Nanosight LM10 (405nm laser, 60mW, software version 3.00064). Samples were analyzed for 60 seconds (count range of 20-100 particles per frame). All measurements were done in triplicate. Alternatively, NTA performed on a ZetaView (Particle Metrix), as previously described^45^

### Atomic Force Microscopy

The EVs were analyzed and characterized by atomic force microscopy (AFM). For the preparation of the samples, the isolated EVs were diluted at 1:20 in ultrapure water and AFM measurements were performed in a BioScope Catalyst atomic force microscope (Bruker), as previously reported^45^.

### EV Protein Extraction and Digestion

To prepare EVs for LC-MS/MS, ∼25μg protein quantified by BCA were lyophilized to dryness and reconstituted in 8M Urea, 50mM ammonium bicarbonate (ABC), 10mM dithiothreitol (DTT), 2% SDS lysis buffer. EV proteins were sonicated with a probe sonicator (3 X 0.5s pulses; Level 1) (Fisher Scientific, Waltham, MA), reduced in 10mM DTT for 30 minutes at room temperature (RT), alkylated in 100mM iodoacetamide for 30 minutes at RT in the dark, and precipitated in chloroform/methanol. On-pellet in-solution protein digestion was performed in 100µL 50mM ABC (pH 8) by adding Trypsin/LysC (Promega, 1:50 ratio) to precipitated EV proteins. EV proteins were incubated at 37°C overnight (∼18h) in a ThermoMixer C (Eppendorf) at 300 rpm. An additional volume of trypsin (Promega, 1:100 ratio) was added for ∼4 hours before acidifying to pH 3-4 with 10% FA.

### SCX Peptide Fractionation and LC-MS/MS

Tryptic peptides were fractionated using strong cation exchange (SCX) StageTips. Briefly, peptides were acidified with 1% TFA and loaded onto a pre-rinsed 12- plug SCX StageTips (Empore™ Supelco, Bellefonte, PA, USA). In total, 6 SCX fractions were collected by eluting in 75, 125, 200, 250, 300 mM ammonium acetate/20% ACN followed by a final elution in 5% ammonium hydroxide/80% ACN. SCX fractions were dried in a SpeedVac (ThermoFisher), re-suspended in ddH2O, and dried again to evaporate residual ammonium acetate. All samples were re-suspended in 0.1% FA prior to LC-MS analysis.

SCX fractions were analyzed using a nanoAquity UHPLC M-class system (Waters) connected to a Q Exactive mass spectrometer (Thermo Scientific) using a nonlinear gradient. Buffer A consisted of water/0.1% FA and Buffer B consisted of ACN/0.1%FA. Peptides (∼1µg estimated by BCA) were initially loaded onto an ACQUITY UPLC M-Class Symmetry C18 Trap Column, 5 µm, 180 µm x 20 mm and trapped for 4 minutes at a flow rate of 5 µl/min at 99% A/1% B. Peptides were separated on an ACQUITY UPLC M-Class Peptide BEH C18 Column (130Å, 1.7µm, 75µm X 250mm) operating at a flow rate of 300 nL/min at 35°C using a non-linear gradient consisting of 1-7% B over 3.5 minutes, 7-19% B over 86.5 minutes and 19-30% B over 30 minutes before increasing to 95% B and washing. Settings for data acquisition on the Q Exactive and Q Exactive Plus are outlined in Supplemental Table 1.

### SCX-DDA Data Analysis

MS raw files were searched in MaxQuant (1.5.2.8) using the Human Uniprot database (reviewed only, updated May 2014 with 40,550 entries). Missed cleavages were set to 3 and I=L. Cysteine carbamidomethylation was set as a fixed modification. Oxidation (M), N-terminal acetylation (protein), and deamidation (NQ) were set as variable modifications (max. number of modifications per peptide = 5) and all other setting were left as default. Precursor mass deviation was left at 20 ppm and 4.5 ppm for first and main search, respectively. Fragment mass deviation was left at 20 ppm. Protein and peptide FDR was set to 0.01 (1%) and the decoy database was set to revert. The match-between-runs feature was utilized across all sample types to maximize proteome coverage and quantitation. Datasets were loaded into Perseus (1.6.14) and proteins identified by site; reverse and potential contaminants were removed47. Protein identifications with quantitative values in >50% samples in each group (cells, plasma or ascites) were retained for downstream analysis unless specified elsewhere. Missing values were imputed using a width of 0.3 and down shift of 1.8 to enable statistical comparisons.

### Label-free Parallel Reaction Monitoring (PRM)

To generate spectral data for biomarker candidate (peptides), several unfractionated plasma EV digests (∼1µg/sample) were initially analyzed on a Q Exactive Plus using a non-linear 2.5h gradient consisting of 1-7% B over 1 minute, 7-23% B over 134 minutes and 23-35% B over 45 minutes before increasing to 95% B and washing. Raw files were searched against the human Uniprot databased (20, 274 entries) using the de novo search engine PEAKS® (version 8). Parent and fragment mass error tolerances were set to 20 ppm and 0.05 Da, respectively. Maximum missed cleavages were set to 3 and 1 non-specific cleavage was allowed. Carbamidomethylation was set as a fixed modification, and deamidation, oxidation and acetylation (protein N-term) were included as variable modifications with a maximum of 3 PTMs per peptide allowed. pepXML peptide information and mzXML spectral data were next exported from PEAKS® generate a PRM method in Skyline^26^. Peptides with missed cleavages or containing tryptophan were removed and up to 3 peptides/protein, 7-18 amino acids in length, were chosen for monitoring. In Skyline, the top 5 most intense transitions (b and y ions) were used for quantification and an 8-minute window was chosen to account for deviations in chromatography and minimize the chance of truncation while maximizing the number of MS/MS scans. EV and EV-depleted samples were subsequently analysed using the same gradient but with a targeted PRM method in a randomized fashion. A minimum of 3 transitions were required to measure peak areas, and targets with dotp scores <0.8 or ppm exceeding 20 were assumed to contain interference and initially assigned a peak area of 0. To correct for sample loading and technical variability, peak areas for each peptide were normalized to the total ion current (TIC). Peak areas were additionally normalized to the CD9 peptide EVQEFYK (extracellular region, AAs 120-126) to correct for EV recovery. Normalized peak areas of 0 were assumed to be missing not at random and imputed with the lowest ratio detected for the given peptide.

### Gas Phase Fractionation Data Independent LC-MS/MS (GPF-DIA)

For spectral library generation, 1µg of plasma EV digest was serially injected to produce 100m/z fractions across 300-1000m/z using a staggered window scheme of 4m/z wide windows that produce 2 m/z bins after demultiplexing, as previously described^37^. 1ug of EV digests from individual donors were analyzed by staggered 24m/z wide windows that produce 12 bins after demultiplexing. Raw files were converted to mzML using ProteoWizard with PeakPicking=1, Demultiplex=10ppm and ZeroSamples=-1. Library and sample mzML files were searched together using DIA-NN to generate a spectral library by allowing 2 missed cleavages and 1 variable modification of Oxidation. Settings for data acquisition on the Eclipse are outlined in Supplemental Table 2.

### Dynamic Retention Time PRM

Using spectral data produced by GPF-DIA and DIA-NN, peptide candidates were selected in Skyline (v23.0.9) by filtering for precursors with CV <30% and minimum product ions of 3. EV digests from the early-stage donors cohort were analyzed using real-time retention time calibrated PRM. Samples were spiked with 50fmol of Pierce Retention Time Calibration (PRTC; ThermoFisher Scientific) mix that allowed for the curation of retention time windows of 3-minutes and selection of peptides with a correlation >0.95 between observation and predicted iRT. Furthermore, we employed the “Dynamic Retention Time” feature in XCalibur software to monitor chromatography shifts in PRTC peptides that allow for real-time correction of downstream PRM windows. In Skyline, the top 8 most intense transitions (b and y ions) were used for quantification. A minimum of 3 transitions were required to measure peak areas, and targets with dotp scores <0.4 were assumed to contain interference and initially assigned a peak area of 0. To correct for sample loading and technical variability, peak areas for each peptide were normalized to the TIC or PRTC. Normalized peak areas of 0 were assumed to be missing not at random and imputed with the lowest ratio detected for the given peptide.

### Detection of CA15-3 Antigen using ELISA

Plasma samples were diluted 2-fold using ddH20 prior to a 30- fold dilution using Dilutent A of the commercial ELISA kit (ThermoFisher Scientific). A standard curve ranging from 1000 U/mL to 4.1 U/mL was used to determine the concentration of MUC1/CA15-3 by analyzing HRP at 450nm and 570nm for background correction using a Cytation (v3.11.19).

### Machine Learning and Statistical Analyses

Differential protein abundance between conditions were determined using a two-tailed Welch’s t-test (p<0.05) in Perseus (version 1.6.14). Graphing was performed using Python or Prism version 6.01 (GraphPad Software, San Diego, CA). Mann-Whitney rank sum statistical tests were calculated in R (version 3.60) or Python (version 3.10). Data handling and machine learning optimization pipelines were built in Python (version 3.10). Pathway and annotation enrichment analyses were performed using Metascape (metascape.org) using the default settings.

## Acknowledgements

We thank Paula Pittock for technical support with UPLC-MS/MS. We would like to thank Dr. Sheela Abraham for access to NTA equipment. This work was supported by the Sawin-Bladwin Chair in Ovarian Cancer Research and the Dr. Anthony Noujaim Oncology Chair awarded to LMP by the Women and Children Health Research Institute and the Alberta Cancer Foundation, respectively. LMP, GAL, and TTC were awarded an operating grant by Cancer Research Society for experimentation with samples obtained from the CR-CHUM biobank.

## Author Contributions

TTC, DDC, and LMP designed the research and wrote the manuscript. TTC, DDC, JL, GMS, LV, and DP conducted experiments. TTC, DDC, JL, GMS, LV, and DP analyzed data and interpreted the results. JDL, GAL, FL and LMP provided logistic and financial support for experimental work. GAL and LMP supervised the study.

