## Supplemental Figure and Tables for "Targeted Proteomics of Plasma Extracellular Vesicles Uncovers MUC1 as Combinatorial Biomarker for the Early Detection of High-grade Serous Ovarian Cancer"

Date: March 1st, 2024

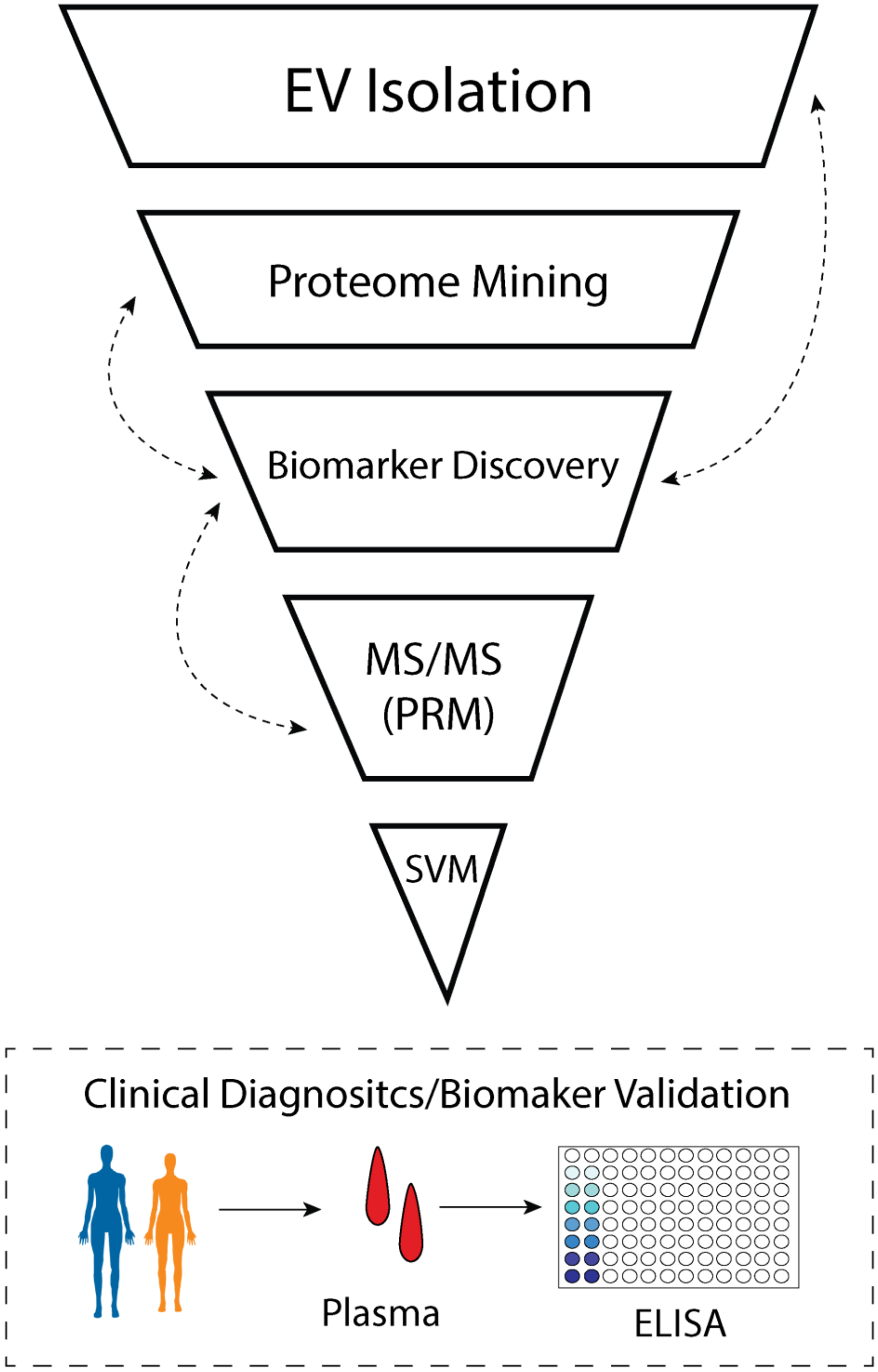

*Graphical Abstract*

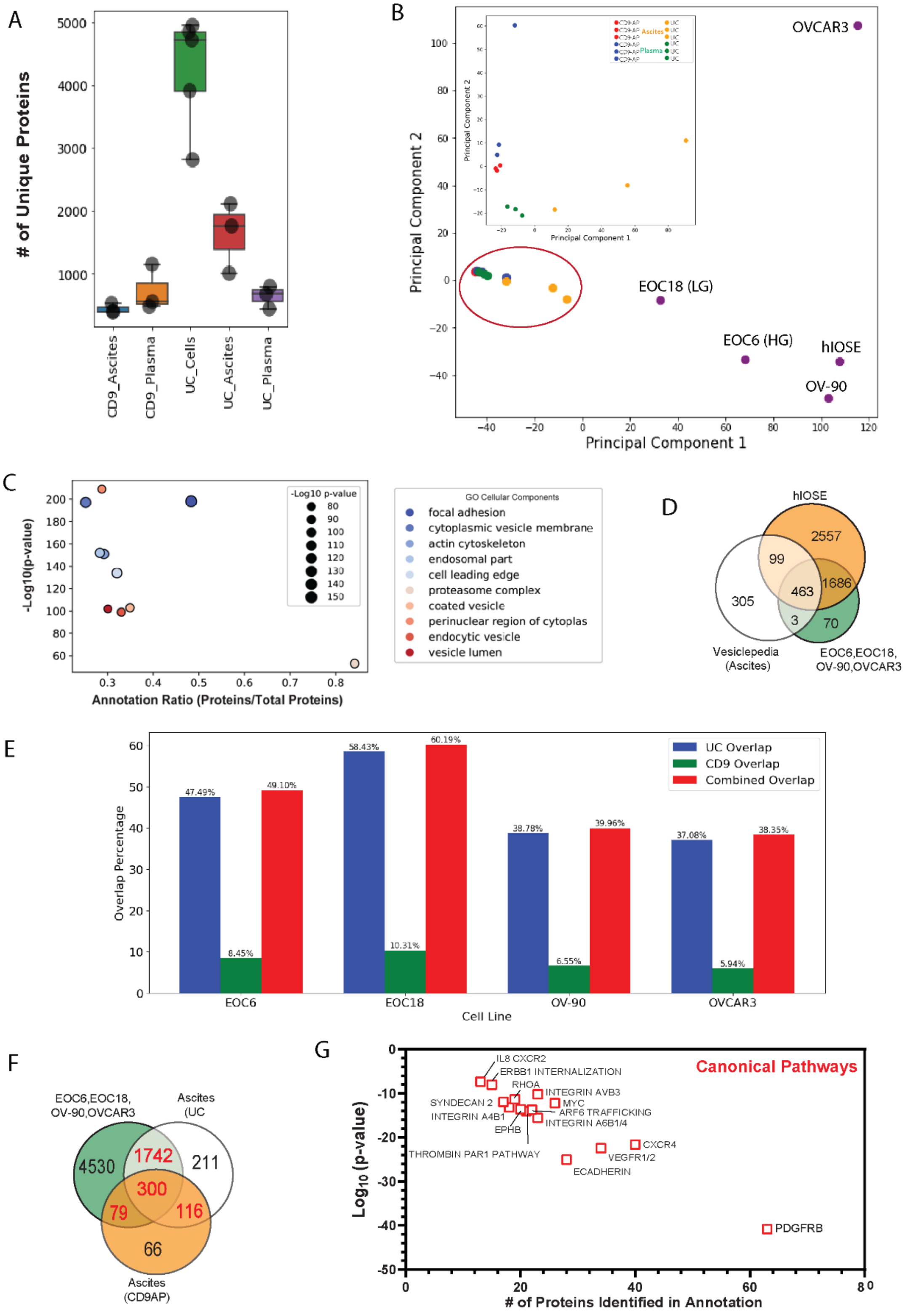

*Supplemental Figure 1*

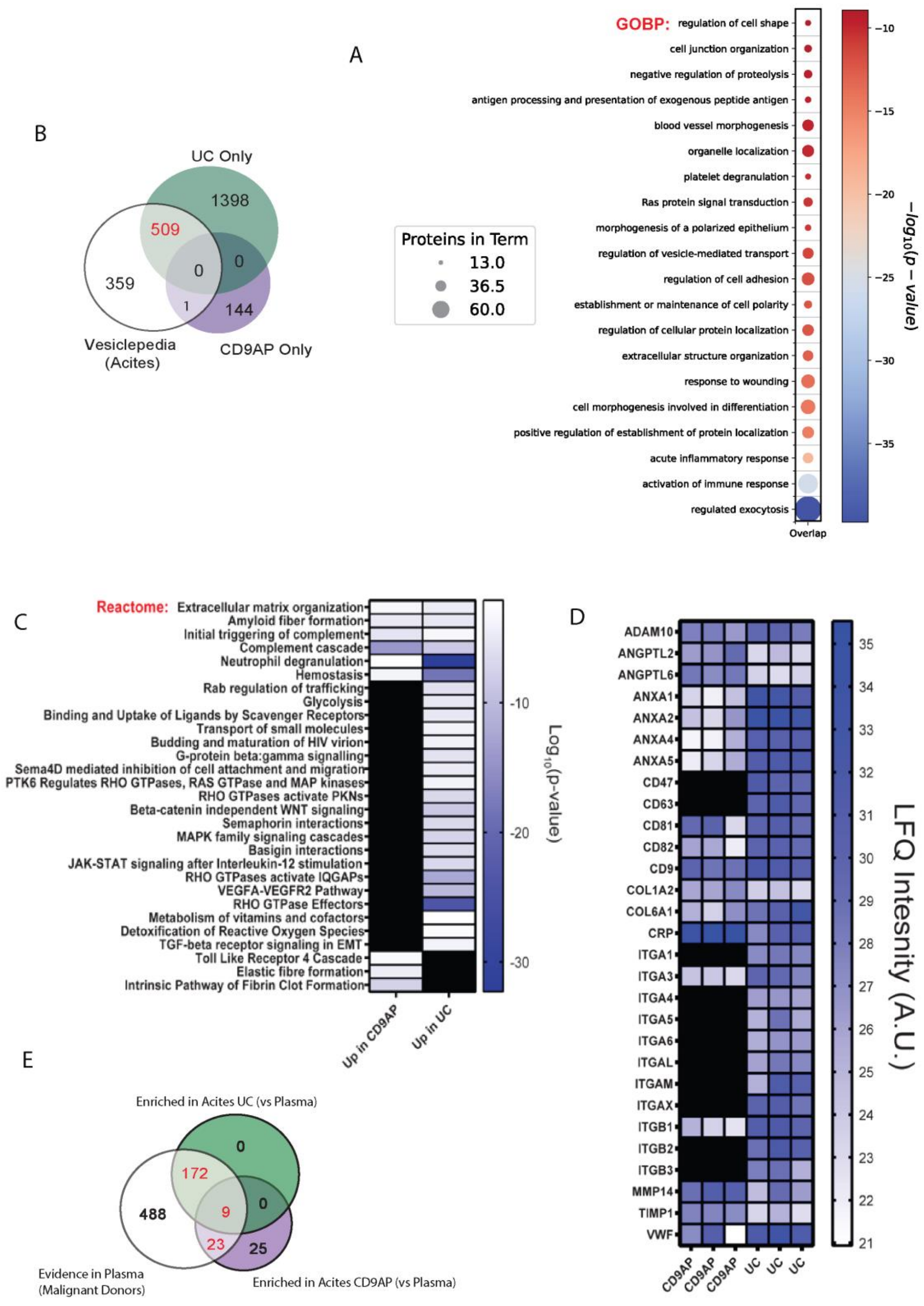

*Supplemental Figure 2*

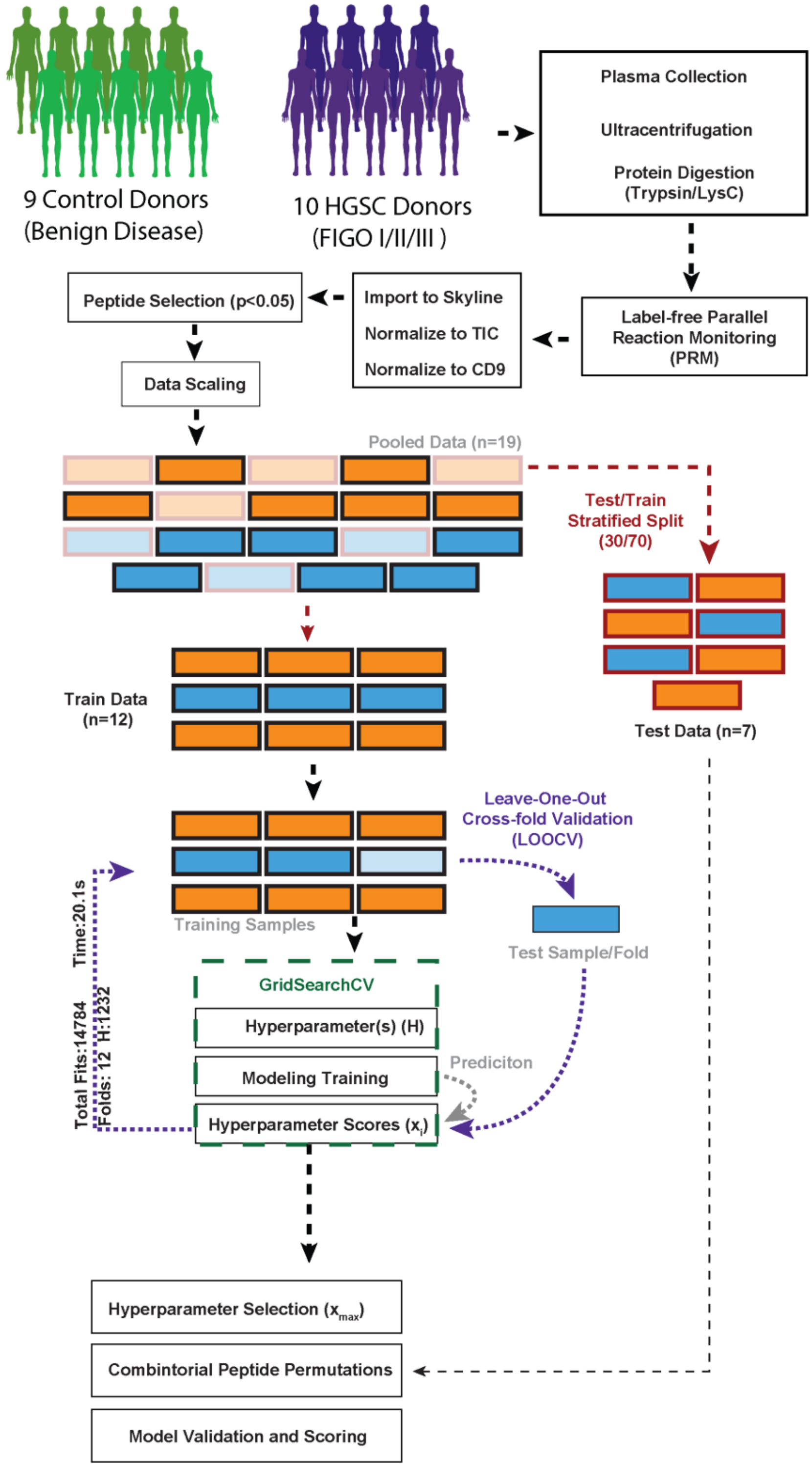

*Supplemental Figure 3*

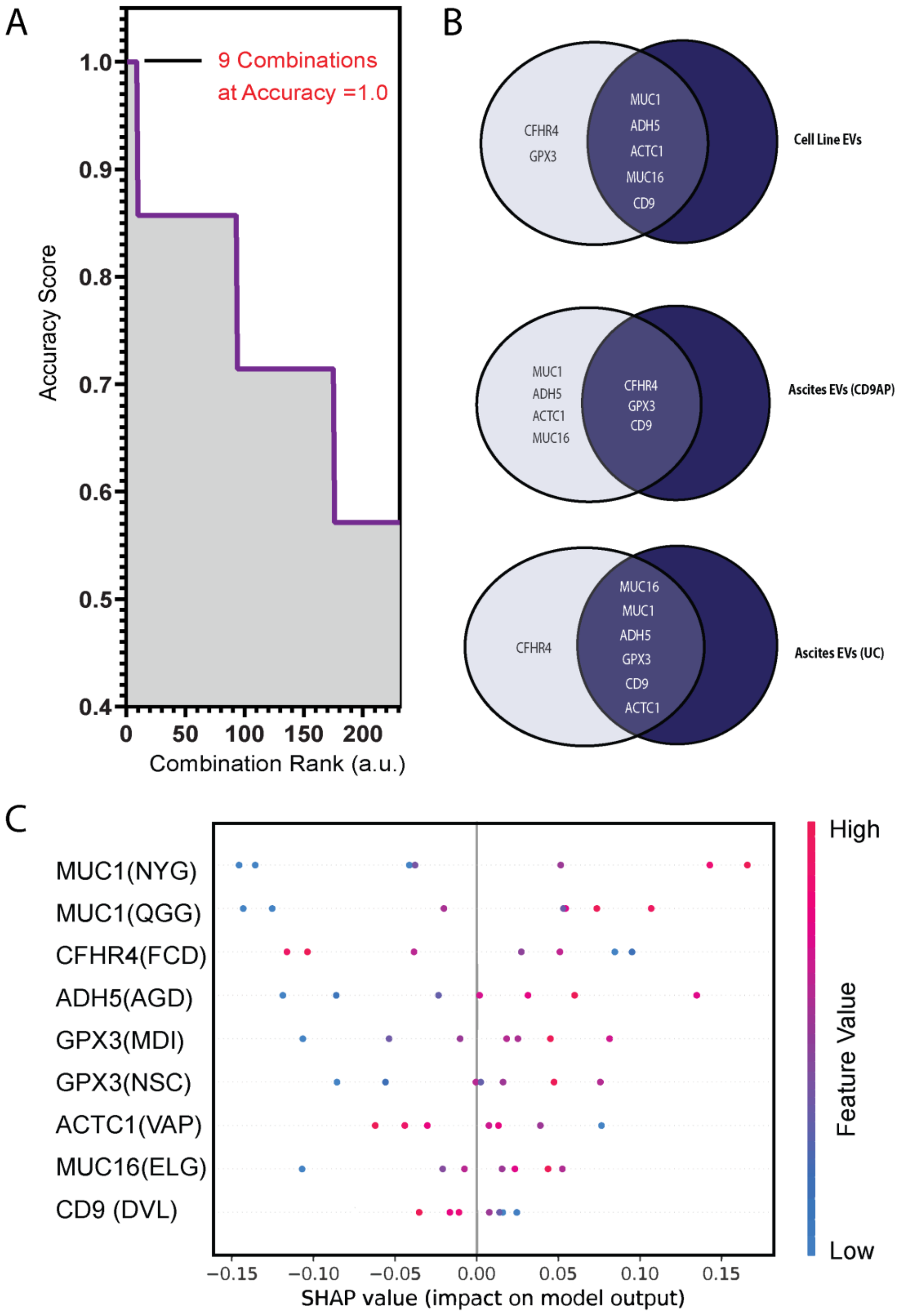

*Supplemental Figure 4*

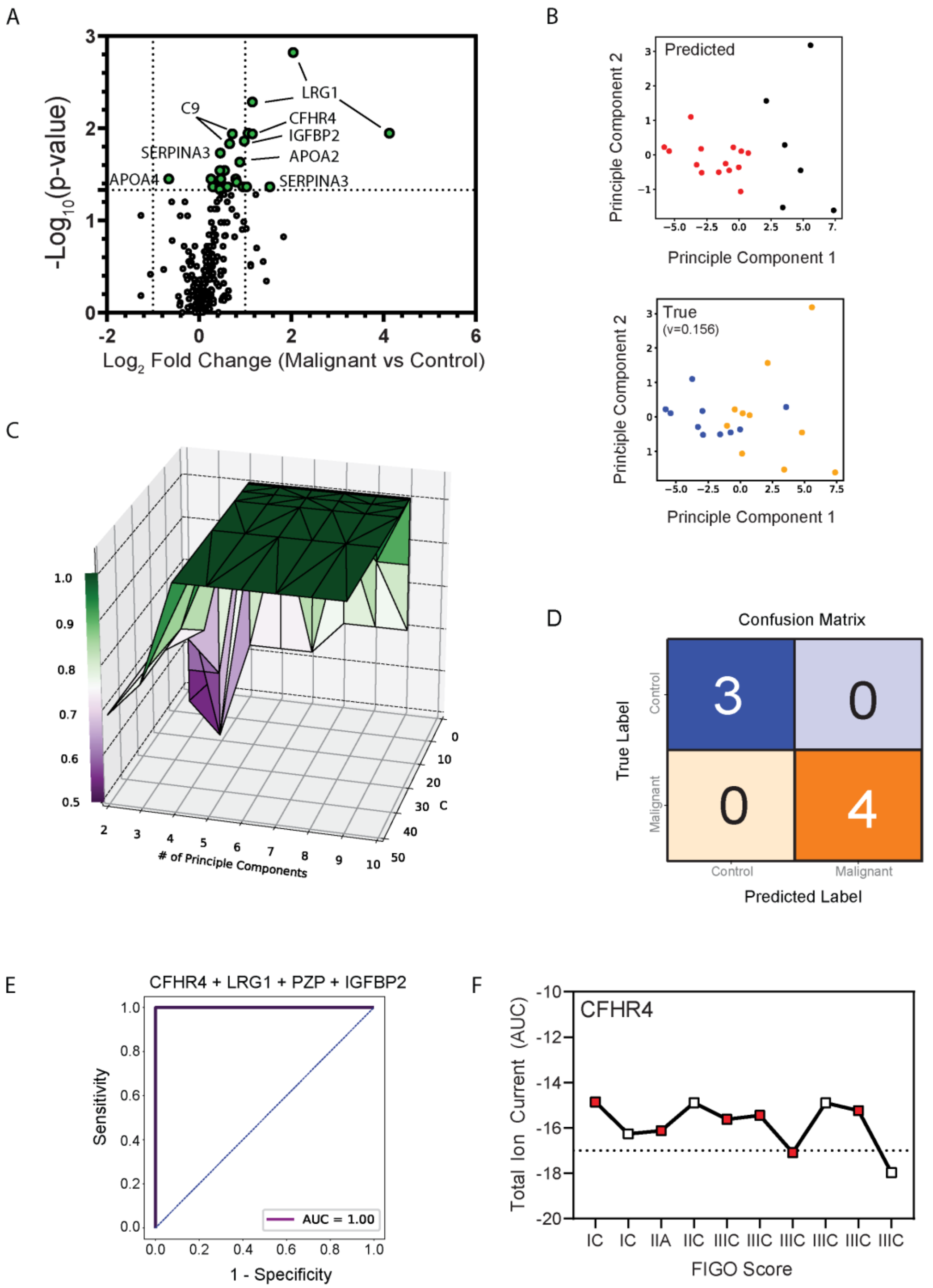

*Supplemental Figure 5*

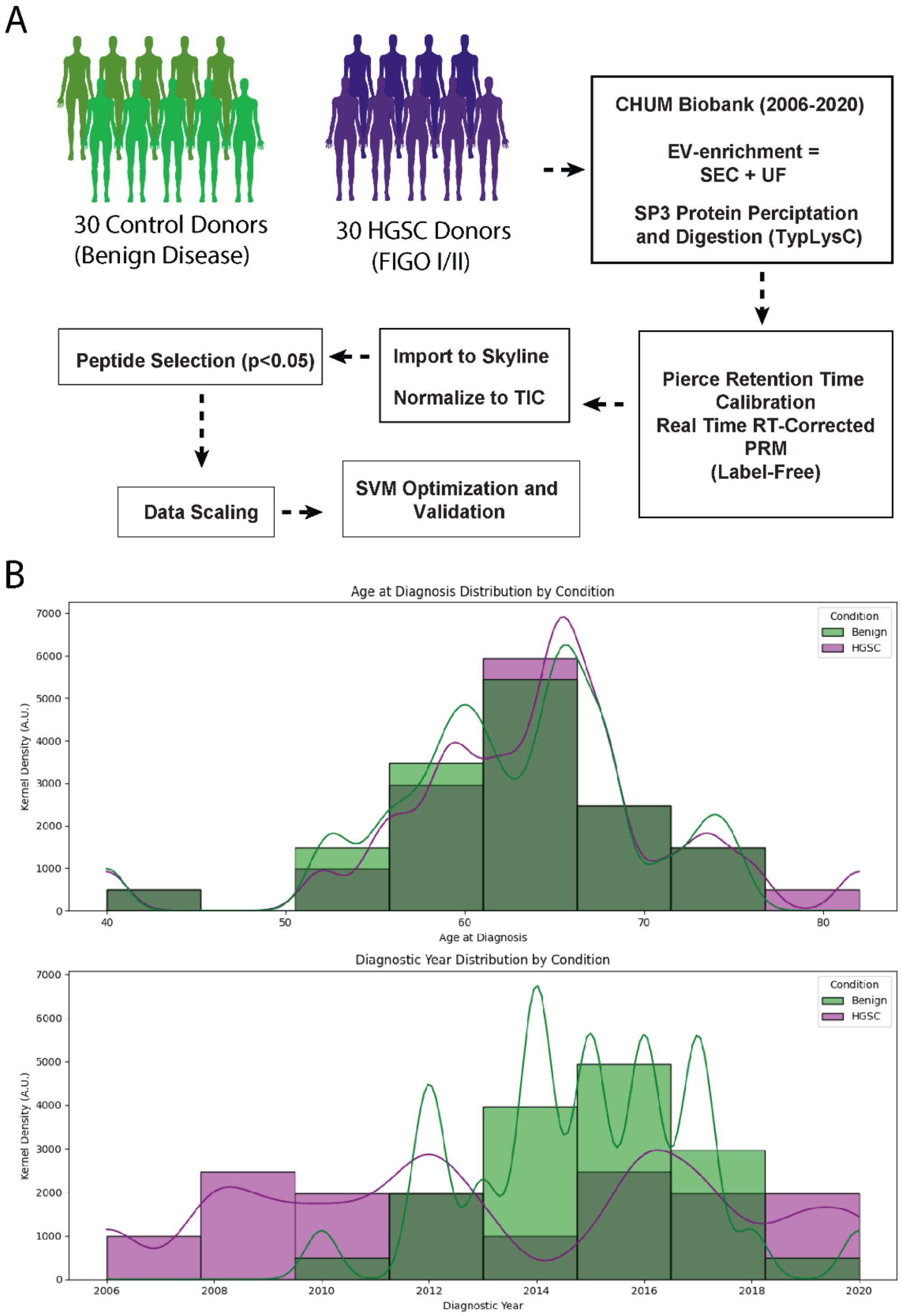

*Supplemental Figure 6*

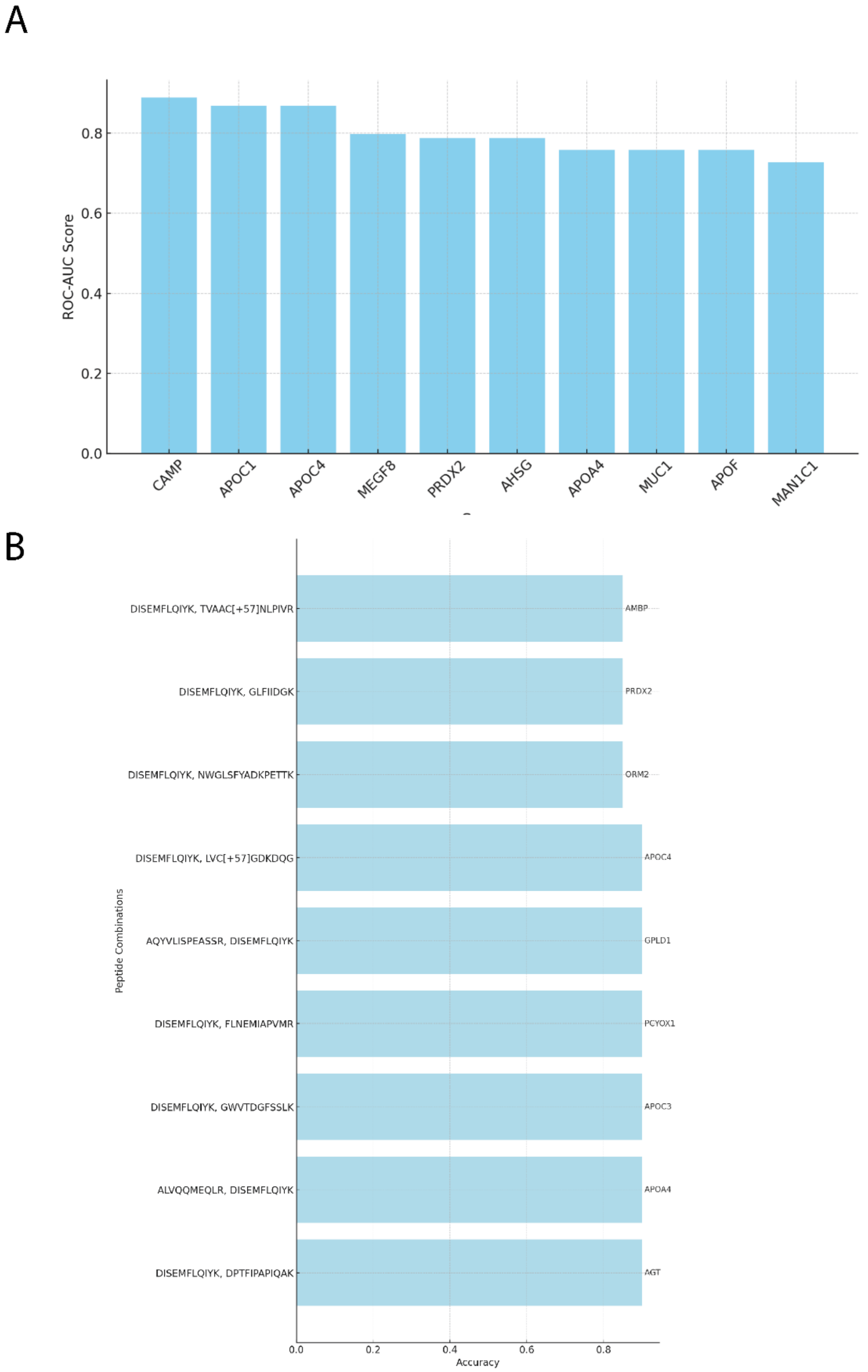

*Supplemental Figure 7*

**Supplemental Table 1.** Q Exactive (Plus) instrument parameters for data acquisition.

| Parameter | Q Exactive | Q Exactive Plus | Q Exactive Plus (PRM settings) |
|---|---|---|---|
| **Orbitrap resolution (MS1)** | 70K | 70K | 70K |
| **Mass range** | 400-1500m/z | 400-1500m/z | 395-1500m/z |
| **MS1 injection time** | 250ms | 250ms | 250ms |
| **MS1 AGC taget** | 3E+06 | 3E+06 | 3E+06 |
| **Lock mass** | 445.120025 | 445.120025 | 445.120025 |
| **MS2 detection** | FT | FT | FT |
| **MS2 resolution** | 17.5K | 17.5K | 35K |
| **MS2 AGC target** | 2E+05 | 2E+05 | 1E+06 |
| **MS2 injection time** | 64ms | 64ms | 120ms |
| **Loop count** | 12 | 12 | 30 |
| **Isolation width** | 1.2 | 1.2 | 1.2 |
| **Isolation offset** | 0.5m/z | 0.5m/z | 0.5m/z |
| **MS2 Activation** | HCD | HCD | HCD |
| **Normalized Collision Energy** | 25 | 25 | 25 |
| **Dynamic exclusion** | enabled | enabled | n/a |
| **Minimum AGC target** | 2.0E+03 | 2.0E+03 | n/a |
| **MS2 intensity threshold** | 3.1E+04 | 3.1E+04 | n/a |
| **Exclusion duration** | 30s | 30s | n/a |
| **Charge exclusion** | unassigned, 1, 7, >8 | unassigned, 1, 7, >8 | n/a |

**Supplemental Table 2. Eclipse Instrument Settings for GPF-DIA Library, GPF-DIA Acquisition, and PRM.**

| PARAMETER | GPF-DIA LIBRARY | GPF-DIA SAMPLE | PRM |
|---|---|---|---|
| MS1 RESOLUTION | **60K** | **60K** | **120K** |
| MS1 SCAN RANGE | **400-1000** | **400-1000** | **400-1000** |
| MS1 ACG TARGET | **100** | **100** | **100** |
| MS1 MAX IT | **50** | **50** | **Auto** |
| MS2 RESOLUTION | **30K** | **30K** | **60K** |
| MS2 SCAN RANGE | **Auto** | **Auto** | **200-1800** |
| MS2 ACG TARGET | **200** | **200** | **200** |
| ISO. WINDOW (M/Z) | **50 x 4 m/z (50%)** | **50 x 24 m/z (50%)** | **1.2** |
| NCE | **33** | **33** | **33** |
| MS2 MAX IT | **54** | **54** | **118** |
| DYNAMIC RT | **N/A** | **N/A** | **Pierce PRTC Mixture** |

**Supplemental Table 3.** Nanoparticle Tracking Analysis on Ascites EVs isolated by Ultracentrifugation or CD9 Immunopurification.

| Parameter | UC-EVs | CD9AP-EVs |
|---|---|---|
| Concentration (particles/mL) | $4.32 \times 10^8 \pm 3.74 \times 10^7$ | $6.07 \times 10^8 \pm 1.37 \times 10^8$ |
| Particles/frame | 21.9 ± 1.9 | 33.4 ± 7.5 |
| Completed tracks (Sum) | 8042 | 16057 |
| Mean | 266.6 ± 14.8 | 106.4 ± 10.5 |
| Mode | 183.5 ± 13.5 | 81.3 ± 3 |
| D10 | 153.7 ± 9.9 | 62.1 ± 1.5 |
| D50 | 204.1 ± 15.1 | 85.5 ± 5.8 |
| D90 | 519.4 ± 5.8 | 166.2 ± 27.6 |

**Supplemental Table 4.** Patient characteristic of malignant PRM samples

| | |
|---|---|
| **Number of samples** | 10 |
| **Age at diagnosis** | |
| **Mean** | 54.8 |
| **Median** | 54 |
| **Range** | 39-69 |
| **FIGO stage** | |
| **IC** | 2 |
| **IIA** | 1 |
| **IIC** | 1 |
| **IIIC** | 6 |
| **Alive** | 5 |
| **Deceased** | 5 |

**Supplemental Table 5. EV-enriched Blood Plasma Peptides Selected for Targeted Proteomics and SVM model optimization.**

| Peptide | Gene | $Log_2$ Fold Change (M vs. C) | $-Log_{10}$ (p-value) | SVM Model Relevance (% of SVM Model) |
|---|---|---|---|---|
| MDILSYMR | GPX3 | 1.31 | 2.25 | 33% |
| QGGFLGLSNIK | MUC1 | 2.57 | 2.03 | 33% |
| FCDMPVFENSR | CHFR4 | 1.06 | 1.89 | 33% |
| YVPPSSTDR | MUC1 | 3.11 | 2.70 | 22% |
| AGDTVIPLYIPQCGECK | ADH5 | -1.20 | 1.40 | 22% |
| DVLETFTVK | CD9 | -0.24 | 1.34 | 22% |
| NSCPPTSELLGTSDR | GPX3 | 0.98 | 1.61 | 11% |
| ELGPYTLDR | MUC16 | 2.12 | 1.22 | 11% |
| VAPEEHPTLLTEAPLNPK | ACTC1 | -0.98 | 2.21 | 0% |
| NYGQLDIFPAR | MUC1 | 1.98 | 2.08 | 0% |
| IISIMDEK | PZP | 1.80 | 2.04 | 0% |
| AYAAGFGDR | TNC | 1.74 | 1.76 | 0% |
| LDAPSQIEVK | TNC | 1.61 | 1.76 | 0% |
| VISQIAMNDEK | SLC34A2 | 2.29 | 1.66 | 0% |
| FEIENCLANK | PZP | 1.47 | 1.66 | 0% |
| ASSFLGEK | C4B | 1.89 | 1.65 | 0% |
| AYSLFSYNTQGR | APCS | 0.89 | 1.56 | 0% |
| EDSPFALK | PZP | 1.66 | 1.55 | 0% |
| ATAQMLEVMFK | LBP | 0.91 | 1.46 | 0% |
| SIPQVSPVR | CPN1 | 0.60 | 1.45 | 0% |
| VATYLPAPEGLK | TNC | 1.61 | 1.36 | 0% |
| DNELLVYK | APCS | 0.83 | 1.34 | 0% |

**M = Malignant, C=Control, SVM= Support Vector Machine, *Linear SVM (C=0.025)**

**Supplemental Table 6.** Patient characteristic of CHUM Samples PRM samples

| | |
|---|---|
| **Number of samples** | 60 |
| **Age at diagnosis** | |
| **Mean** | 62.8 |
| **Median** | 64 |
| **Range** | 40-82 |
| **FIGO stage** | |
| **IA** | 7 |
| **IB** | 1 |
| **IC** | 8 |
| **IIA** | 6 |
| **IIB** | 7 |
| **IIC** | 1 |
